# SUMO Promotes DNA Repair Protein Collaboration to Support Alterative Telomere Lengthening in the Absence of PML

**DOI:** 10.1101/2024.02.29.582813

**Authors:** Rongwei Zhao, Meng Xu, Anne R. Wondisford, Rachel M. Lackner, Jayme Salsman, Graham Dellaire, David M. Chenoweth, Roderick J. O’Sullivan, Xiaolan Zhao, Huaiying Zhang

**Affiliations:** Department of Biology, Carnegie Mellon University, Pittsburgh, PA 15213, USA; Department of Pharmacology and Chemical Biology, UPMC Hillman Cancer Center, University of Pittsburgh, Pittsburgh, PA 15213, USA; Department of Chemistry, University of Pennsylvania, Philadelphia, PA 19014, USA; Department of Pathology, Dalhousie University, Halifax, Nova Scotia, B3H 4R2, Canada; Molecular Biology Program, Memorial Sloan Kettering Cancer Center, New York, NY 10065

## Abstract

Alternative lengthening of telomeres (ALT) pathway maintains telomeres in a significant fraction of cancers associated with poor clinical outcomes. A better understanding of ALT mechanisms can provide a basis for developing new treatment strategies for ALT cancers. SUMO modification of telomere proteins plays a critical role in the formation of ALT telomere-associated PML bodies (APBs), where telomeres are clustered and DNA repair proteins are enriched to promote homology-directed telomere DNA synthesis in ALT. However, whether and how SUMO contributes to ALT beyond APB formation remains elusive. Here, we report that SUMO promotes collaboration among DNA repair proteins to achieve APB-independent telomere maintenance. By using ALT cancer cells with PML protein knocked out and thus devoid of APBs, we show that sumoylation is required for manifesting ALT features, including telomere clustering and telomeric DNA synthesis, independent of PML and APBs. Further, small molecule-induced telomere targeting of SUMO produces signatures of phase separation and ALT features in PML null cells in a manner depending on both sumoylation and SUMO interaction with SUMO interaction motifs (SIMs). Mechanistically, SUMO-induced effects are linked to the enrichment of DNA repair proteins, including Rad52, Rad51AP1, and BLM, to the SUMO-containing telomere foci. Finally, we find that Rad52 can undergo phase separation, enrich SUMO on telomeres, and promote telomere DNA synthesis in collaboration with the BLM helicase in a SUMO-dependent manner. Collectively, our findings suggest that, in addition to forming APBs, SUMO also promotes collaboration among DNA repair proteins to support telomere maintenance in ALT cells. Given the promising effects of sumoylation inhibitors in cancer treatment, our findings suggest their potential use in perturbing telomere maintenance in ALT cancer cells.

## Introduction

To sustain continuous proliferation, cancer cells must maintain their telomeres either by telomerase reactivation or by alternative telomere lengthening (ALT)(Henson et al., 2002; Varley et al., 2002). An estimated 10-15% of cancer types employ ALT and these are often associated with poor survival outcomes(Dilley & Greenberg, 2015; Yeager et al., 1999). Past studies have established that telomere synthesis in ALT is achieved by break-induced replication (BIR), a homologous recombination (HR) mechanism(Cho et al., 2014; Dilley et al., 2016). ALT can either utilize a BIR pathway that depends on the recombination protein Rad52(Min et al., 2017; J. M. Zhang et al., 2019) or another one that requires the repair protein Rad51AP1(Barroso-González et al., 2019; Yadav et al., 2022; J. M. Zhang et al., 2019). Among additional HR proteins involved in ALT, the BLM helicase plays a prominent role by enabling both of the BIR pathways(J. M. Zhang et al., 2019; J.-M. Zhang et al., 2021).

In addition to HR proteins, ALT also relies on sumoylation that conjugates the small ubiquitin modifier (SUMO) to target proteins(Seeler & Dejean, 2003). Sumoylation of telomere proteins, such as TRF1 and TRF2, can promote the formation of ALT-specific nuclear bodies called ALT-associated PML bodies (APBs)(Brouwer et al., 2009; Chung et al., 2011; Potts & Yu, 2007). APB formation also depends on the PML protein, which is both sumoylated and contains SUMO interaction motifs (SIMs)(Kamitani et al., 1998; Shen et al., 2006). The multi-valent SUMO-SIM interactions mediated by PML and telomere proteins enable APB formation via phase separation(H. Zhang et al., 2020). It is thought that APBs facilitate ALT by enriching telomere clusters (Draskovic et al., 2009; Heaphy et al., 2011) and DNA repair proteins within the same space(Dilley et al., 2016; Roumelioti et al., 2016). Despite the critical roles of APBs in ALT, a recent study examined cancer cell lines that rely on ALT for telomere maintenance and found that while PML protein knockout abolished APBs and reduced ALT, cells are viable for months(Loe et al., 2020). This finding suggests the possibility of APB-independent telomere lengthening in PML null cells. However, the nature of this process and the role of sumoylation in ALT beyond APB formation remain elusive.

Here, we address the above questions by examining PML null ALT-positive U2OS cells lacking APBs. We found SUMO is enriched at telomeres even in the absence of PML and APBs. Importantly, telomere breaks induced by the FokI nuclease fused to the telomere protein TRF1 can induce two key ALT features in PML null cells, namely telomere clustering and telomeric DNA synthesis. This provides evidence that ALT can take place in the absence of PML and APBs. We further applied two experimental strategies not previously used in ALT studies. These include the application of sumoylation inhibitors and chemical-induced protein targeting, which allows live cell imaging and prevents toxicity caused by constitutive targeting. Results derived from SUMO inhibitor studies revealed that sumoylation is required for ALT features induced by telomere DNA cleavage by FokI in PML null cells. This result strongly argues that sumoylation can contribute to ALT in an APB-independent manner. Data from the chemical-induced protein telomere targeting technique showed that transiently targeting SUMO to telomeres induces ALT features independent of PML and this requires both sumoylation and SUMO-SIM interaction.

Finally, we show that targeting SUMO or Rad52 to telomeres mutually enriches each other at telomers and both induce signatures of phase separation and promote telomere synthesis in PML null cells. Collectively, our data provided several lines of evidence to support the conclusion that sumoylation and DNA repair proteins collaborate to support ALT features beyond promoting APB formation.

## Results

### Sumoylation contributes to ALT features even in the absence of PML and APBs

To address whether SUMOylation contributes to ALT independent of APBs and PML, we utilized PML knockout (PML KO) U2OS cells devoid of APBs(Loe et al., 2020). We confirmed the absence of nuclear bodies containing PML in these cells using immunofluorescent staining (Fig. S1A). As seen previously (Loe et al., 2020), the growth of U2OS cells was largely unaffected by the loss of PML despite impairment in ALT. These cells allowed us to address possible roles of SUMO in ALT that are independent of APBs and PML.

ALT features can be induced by telomere-specific DNA breaks generated by the FokI nuclease targeted to telomeres through fusing to the telomeric protein TRF1(Cho et al., 2014; Dilley et al., 2016). While previous studies used this system in PML-containing cells, we employed it in PML null U2OS cells to address whether telomere breaks can induce ALT features without PML and APBs and whether these features require SUMO. First, we assessed whether SUMO was localized to telomeres after FokI induction. Immunofluorescent imaging revealed that the TRF1-Fok1 fusion, but not the nuclease dead mutant fusion TRF1-FokI-D450A, increased SUMO1 and SUMO2/3 (SUMO2 and SUMO3 detected by the same antibody) localization at telomeres in PML null cells (Fig. 1A, B, Fig. S1B, C). The level of telomeric SUMO in PML null cells was similar to that of control PML-containing cells (Fig. 1A, B). In addition, MMS21 and PIAS4, two SUMO E3 ligases important for ALT(Potts & Yu, 2005, 2007; J.-M. Zhang et al., 2021), were found at telomeres in PML null cells, just like in PML-containing cells (Fig. S1E, F). These data suggest that SUMO and SUMO E3 enrichment at ALT telomeres can occur in response to telomere breaks without PML and APBs.

**Fig. 1.**
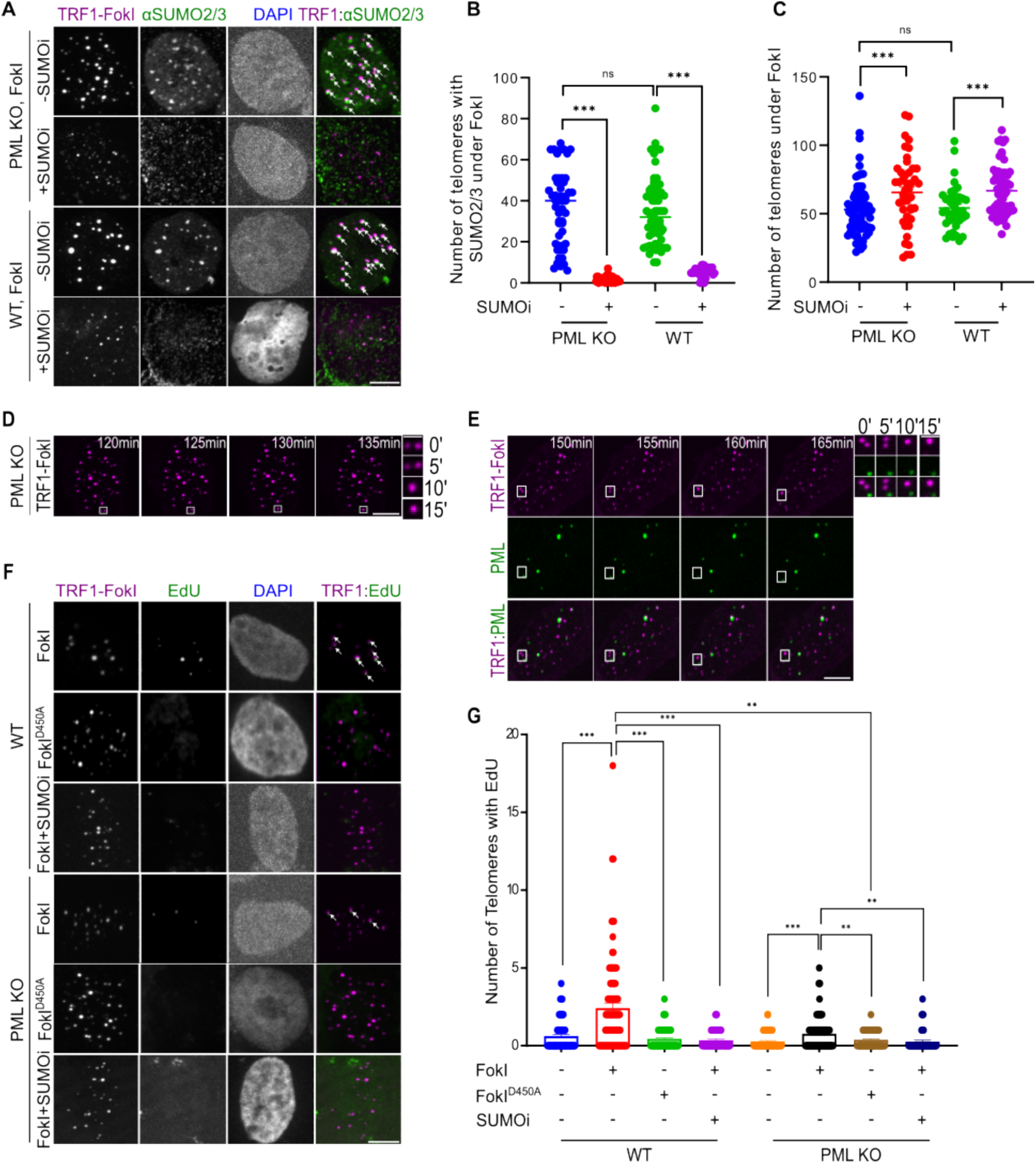
SUMO pathway is required for ALT features regardless of APB status. **(A)** Representative images and **(B)** quantification of SUMO2/3 localization on telomeres and **(C)** telomere numbers in PML knockout (PML KO) U2OS cells expressing mCh-TRF1-FokI with the treatment of 4-Hydroxyestradiol (4-OHT) for 6 hr to translocate FokI to the nucleus with or without 1 μM SUMO inhibitor (SUMOi) for 2 days. (each dot represents one cell, three independent experiments). White arrows indicate SUMO2/3 co-localizations on telomeres. **(D)** Live cell imaging of PML KO cells expressing mCh-TRF1-FokI after inducing DNA damage. **(E)** Representative images of PML-Clover cells expressing mCh-TRF1-FokI after adding 4-OHT at the indicated time. Zoomed-in images on the right side of (D) and (E) show telomere clustering events. **(F)** Representative images and **(G)** quantification of Edu staining for newly synthesized telomeric DNA after 6-hour FokI induction with or without SUMOi for 2 days in U2OS WT and PML KO cells. White arrows indicate EdU signals on telomeres. Scale bars: 1 μm for the zoomed images and 5 μm for other images.

Significantly, TRF1-FokI, but not TRF1-FokI-D450A, induced telomere clustering in PML null cells, evidenced by telomere DNA FISH data that showed a reduction in telomere numbers (Fig. S1D). The degree of telomere clustering in the PML null cells was similar to that in the PML-containing control cells (Fig. 1A, C). Live cell imaging further captured the process of telomere clustering after FokI induction in PML null cells (Fig. 1D, Movie 1), as reported in PML-containing cells(Cho et al., 2014). To test if damaged telomeres can cluster without APBs even in the PML-containing ALT cells, we used a U2OS cell line where endogenous PML is labeled with Clover(Pinder et al., 2015). Live imaging revealed that upon inducing TRF1-FokI, clustered telomeres form independent of forming APBs. (Fig. 1E, Movie 2). Collectively, these data show that telomere breaks can induce telomere clustering without APBs.

Next, we measured nascent telomeric DNA synthesis based on EdU incorporation at telomeres. We observed the induction of telomeric DNA synthesis in PML null cells after FokI induction (Fig. 1F, G). Unlike SUMO localization (Fig. 1B) and telomere clustering (Fig. 1C), telomeric DNA synthesis in PML null cells was less pronounced than that in PML-containing control cells, suggesting that APBs are more important for telomeric DNA synthesis than SUMO enrichment and telomere clustering. In conclusion, while our data are consistent with an established role of PML/APBs in telomere DNA synthesis in ALT cells(O’Sullivan et al., 2014; Sahin Umut et al., 2014), they unveil that in the absence of PML, telomere breaks can induce three key ALT features, namely localization of SUMO to telomeres, telomere clustering, and telomeric DNA synthesis.

Finally, we asked whether sumoylation is required for telomere clustering and telomere DNA synthesis independent of PML and APBs. To this end, we used a well-established small molecule inhibitor (ML-792) of the SUMO-activating enzyme to downregulate sumoylation(He et al., 2017). Treatment with this SUMO inhibitor (SUMOi) decreased SUMO1 and SUMO2/3 levels at telomeres after TRF1-FokI expression in PML null and PML-containing cells (Fig. 1A, B, Fig. S1G, H), confirming the effectiveness of the inhibitor. Significantly, SUMOi treatment decreased telomere clustering upon TRF1-FokI induction in both types of cells (Fig. 1C). In addition, in both PML null and PML-containing cells, SUMOi treatment decreased telomeric DNA synthesis upon TRF1-FokI induction to a level similar to that seen either without FokI expression or expressing the FokI mutant TRF1-FokI-D450A (Fig. 1F, G). These results suggest that upon induction of telomere breaks, sumoylation is required for ALT telomere clustering and telomere DNA synthesis in the absence of PML/APBs.

### Sumoylation is required for endogenous telomere clustering independent of APBs

We next examined how endogenous ALT is affected by sumoylation in the absence of APBs. First, we arrested the PML null cells in the G2 phase when ALT was active and assessed whether SUMO was localized to telomeres. Immunofluorescent imaging revealed that SUMO1 and SUMO2/3 are localized to telomeres at a level moderately lower than those seen in the control cells with intact PML (Fig. S2A-D). This result suggests that SUMO has an intrinsic ability to be enriched at ALT telomeres, and this can be enhanced by PML in the endogenous ALT pathway.

Next, we asked whether SUMO inhibitors affected ALT features in G2-arrested PML null cells. SUMOi decreased SUMO1 and SUMO2/3 levels at telomeres both in PML null and PML-containing cells (Fig. S2A-D). Significantly, SUMOi treatment decreased telomere clustering in both types of cells (Fig. S2E). This is in addition to reduced numbers of APBs and PML bodies seen in cells containing PML, as expected from an established role of SUMO in the formation of PML bodies(Hirano & Udagawa, 2022; Zhong et al., 2000) and APBs (Chung et al., 2011; Potts & Yu, 2007) (Fig. S2F, G). These data reveal an APB-independent role of sumoylation during endogenous ALT.

### Targeting SUMO to telomeres induces signatures of phase separation and ALT features in PML null cells

We have previously shown that in PML-containing cells, SUMO-SIM interactions on telomeres induce telomere clustering and signatures of phase separation(H. Zhang et al., 2020). We asked whether enriching SUMO at telomeres would have the same effects in PML null cells. To this end, we used an inducible protein dimerization system that can transiently and effectively recruit proteins to specific genomic loci(Lackner et al., 2022; Zhao et al., 2021). In our setup, each SUMO isoform (SUMO1, 2, and 3) was fused to mCherry and the eDHRF protein, while the telomere protein TRF1 was fused to GFP and the 3xHalo-enzyme (Fig. 2A). The addition of the chemical dimerizer, trimethylolpropane-fluorobenzamide-halo ligand (TFH), that binds to both eDHRF and Halo-enzyme can induce interaction between the two fusion proteins. This technique can minimize toxicity associated with constitutive SUMO enrichment at telomeres and is compatible with live cell imaging for the examination of key ALT features in real-time.

**Fig. 2.**
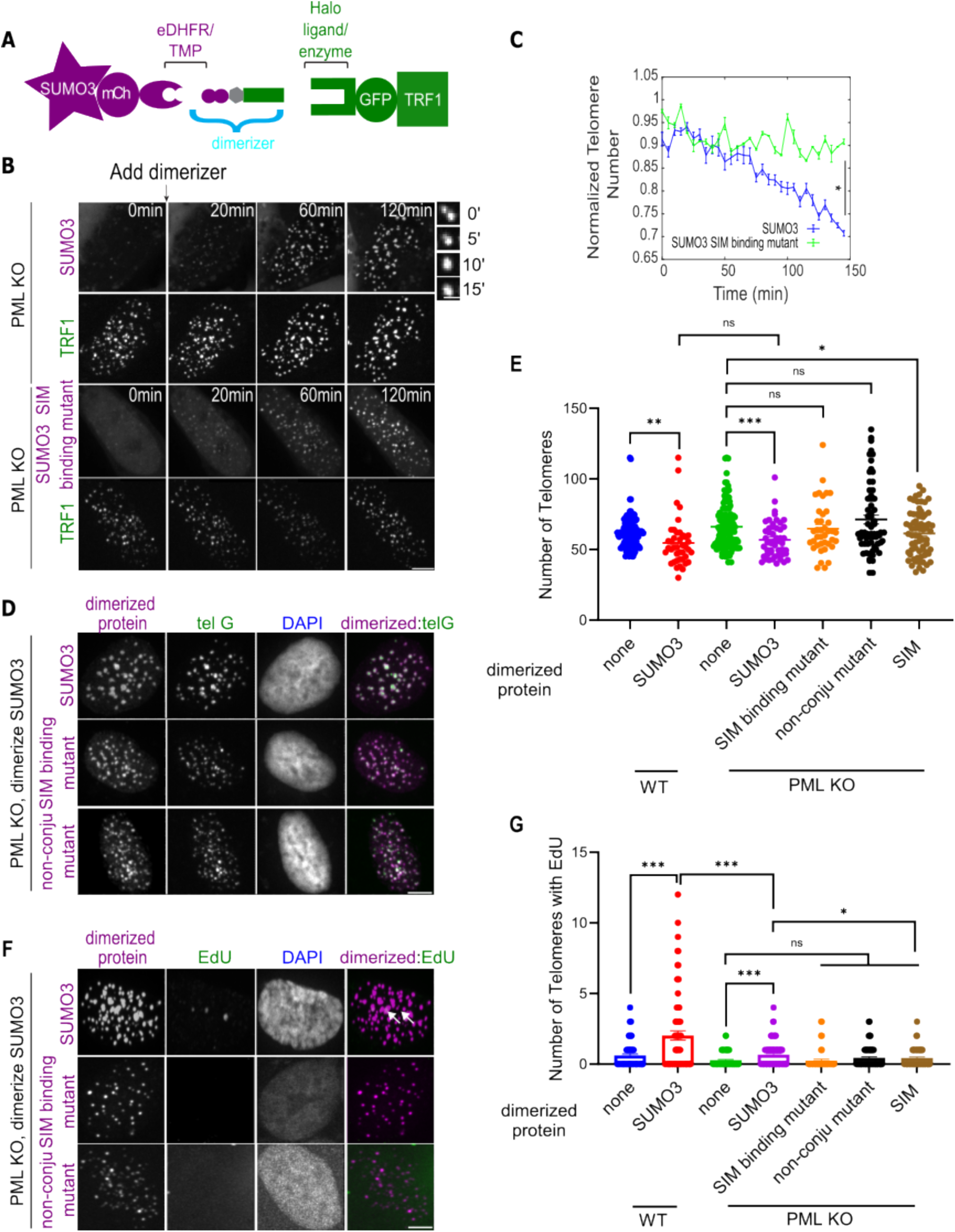
SUMO enrichment induces signatures of phase separation and ALT phenotypes regardless of APBs. **(A)** Dimerization schematic: SUMO3 is fused to mCherry and eDHFR, and TRF1 is fused to GFP and 3xHalo enzyme. The dimerizer, TMP (trimethoprim)-Fluorobenzamide-Halo ligand (TFH), can interact with eDHFR and Halo enzyme to dimerize SUMO3 to TRF1. **(B)** Representative images of PML KO cells after dimerizing mCh-eDHFR-SUMO3 or SUMO3 SIM binding mutant to 3xHalo-GFP-TRF1 after the first time point. Zoomed-in images on the right show the fusion of TRF1 foci after dimerizing SUMO3. **(C)** Telomere number per cell over time from live images after dimerizing SUMO3 or SUMO3 mutant that cannot interact with SIM. Telomere numbers are normalized by the number at the first time point for each cell (more than 22 cells per group, three independent experiments, two-tailed unpaired *t-test)*. **(D)** Representative images of telomere DNA FISH and **(E)** quantification of telomere number after dimerizing SUMO3, SUMO3 SIM interacting mutant, SUMO3 non-conjugable mutant, and SIM to telomeres for 6 hours. (more than 60 cells per group, three independent experiments). **(F)** Representative images and **(G)** quantification of Edu staining for newly synthesized telomere DNA with/without dimerizing SUMO3, SUMO3 mutants, and SIM to telomeres in PML KO cells. Telomeres were visualized by FISH (images not shown). White arrows indicate EdU foci on telomeres. Scale bars: 1 μm for the zoomed images and 5 μm for other images.

Successful recruitment of each SUMO isoform to TRF1-GFP marked telomeres upon the addition of the dimerizer in the PML null cells was confirmed with microscopy (Fig. 2B). Both SUMO-mCherry and TRF1-GFP formed bright and round foci and the two types of foci showed high degrees of colocalization (Fig. 2B, Fig. S3A, Movie 3-5). Further, these foci showed fusion behaviors characteristic of phase-separated condensates (Fig. 2B, Fig. S3A, Movie 3-5).

Moreover, telomeres clustered after dimerizer addition in PML null cells, as reflected in the reduced telomere numbers compared with the condition without dimerizer (Fig. 2C). The increased telomere clustering was also confirmed with FISH (Fig. 2D, E, Fig. S3B). Significantly, live imaging of PML-Clover cells after dimerizing SUMO to telomeres showed APB-independent telomere clustering in addition to APB-dependent clustering (Fig. S3C-G). Finally, recruiting SUMO isoforms to telomeres increased telomere DNA synthesis (Fig. 2F, G, Fig. S3H, I). Collectively, the observed effects of dimerizer-induced SUMO recruitment to telomeres provide evidence for PML and APB-independent formation of phase-separated condensates containing SUMO and clustered telomeres that are capable of telomere DNA synthesis.

For all examined effects described above, the three SUMO isoforms behaved similarly. Recruiting one isoform enriched the others on telomeres. As SUMOi reduced such mutual enrichment (Fig. S4A, B), we conclude that sumoylation underlies this isoform inter-dependancy. Focusing on SUMO3, we found that its recruitment to telomeres resulted in a similar level of telomere clustering in either PML-containing or PML KO cells (Fig. 2E), but more telomere DNA synthesis in the former (Fig. 2G), highlighting a role of APBs in ALT telomere synthesis beyond SUMO enrichment and telomere clustering. Collectively, these results suggest that, in PML-free cells, SUMO enrichment at telomeres can induce signatures of phase separation as well as key ALT features, including telomere clustering and telomeric DNA synthesis.

### Sumoylation and SUMO-SIM interaction underlie the PML-independent role of SUMO in ALT

We moved on to examine the mechanisms underlying SUMO-mediated ALT features in the absence of PML and APBs. Previously, we found that recruiting SIM to telomeres in PML-containing cells using the chemically induced dimerization system described above induced APB formation via phase separation and that this requires its SUMO binding capacity(H. Zhang et al., 2020). We asked here whether SUMO-SIM interaction also contributes to ALT in a PML and APB-independent manner.

First, we examined the consequences of targeting SIM to telomeres using a dimerizer system. We found that recruiting SIM to telomeres in PML null cells led to the enrichment of SUMO isoforms at telomeres (Fig. S4C-E). Second, this system generated TRF1-containing round droplets that fuse among themselves over time, which is suggestive of phase separation behavior (Fig. S4G, H). Third, SIM recruitment to telomeres increased telomere clustering (Fig. S4F, H), though less pronounced than that caused by telomere targeting of SUMO (Fig. 2E). Fourth, SIM recruitment to telomeres was insufficient to induce telomere DNA synthesis in PML null cells (Fig. 2G), similar to what we reported in cells containing PML(H. Zhang et al., 2020), but different from targeting SUMO to telomeres that induced telomere synthesis (Fig. 2F, G). We suspect that the difference could be caused by differences in SUMO-SIM stoichiometries, as this feature affects condensate composition(Banani et al., 2016; Ditlev et al., 2018). Regardless, the above findings support the notion that targeting SIM to telomeres can induce a subset of ALT features.

We moved on to investigate whether the SUMO-SIM interaction is important for ALT features. To this end, we utilized SUMO and SIM mutants that are defective in SUMO-SIM binding and targeted them to telomeres using the dimerization system. First, we found that a SUMO3 variant mutated for its SIM binding site (FKIK mutated to FAAA) (Banani et al., 2016) did not induce signatures of phase separation (Fig, 2B, C, Movie 6), and led to less telomere clustering (Fig. 2D, E) and less telomere DNA synthesis (Fig. 2F, G). Second, recruiting a SIM mutant that cannot interact with SUMO (VIDL mutated to VADA) (Banani et al., 2016) failed to generate TRF1-containing droplets and telomere clustering (Fig. S4G, H). Third, recruiting a SUMO3 mutant that could not be conjugated to substrates (C-terminal di-Gly motif mutated to di-Val) (Banani et al., 2016) resulted in less telomere clustering (Fig. 2D, E) and less telomere DNA synthesis (Fig. 2F, G). This is accompanied by a lack of enrichment of SUMO1 and 2 on telomeres (Fig. S4A, B). These results are consistent with each other and suggest that protein sumoylation and SUMO-SIM interaction contribute to SUMO-induced ALT features in PML null cells.

### Sumoylation and SUMO-SIM interaction promote DNA repair factor enrichment at telomeres independent of APBs

We next addressed how SUMO-SIM interaction can promote ALT features in PML null cells. It is known that SUMO and DNA repair proteins colocalize at damage sites in a PML-independent manner(Claessens et al., 2023) and that many DNA repair proteins are often both sumoylated and contain SIMs(Dhingra & Zhao, 2019; Hu & Parvin, 2014; Psakhye & Jentsch, 2012). We thus reasoned that DNA repair factors that both contribute to ALT and contain SUMOylation sites and/or SIMs could mediate SUMO-SIM-dependent effects on ALT. After surveying the literature, we tested three proteins, each of which contains at least one predicted and/or confirmed sumoylation site and SIM site. These include the BLM helicase(Min et al., 2019), the Rad51AP1(Barroso-González et al., 2019), and Rad52 repair factors(Sacher et al., 2006; Silva et al., 2016; Torres-Rosell et al., 2007; Verma et al., 2019). While Rad52 and Rad51AP1 each control one of the two ALT pathways, BLM contributes to both(J. M. Zhang et al., 2019). Thus the behavior of these proteins can help us understand how the SUMO-SIM interaction affects both ALT pathways.

We found that targeting SUMO3 to telomeres in PML null cells induced enrichment of BLM, Rad51AP1, and Rad52 at telomeres (Fig. 3A, B, Fig. S5A, B). Similar effects were seen when SIM was targeted to telomeres, though to a much lesser degree (Fig. S5D, G, H). The differences here are consistent with the relatively weaker effects of SIM on inducing ALT features described above (Fig. 2E, G). These observations suggested the three ALT proteins can be recruited to telomeres upon increasing SUMO abundance at telomeres in the absence of PML and APBs.

**Fig. 3.**
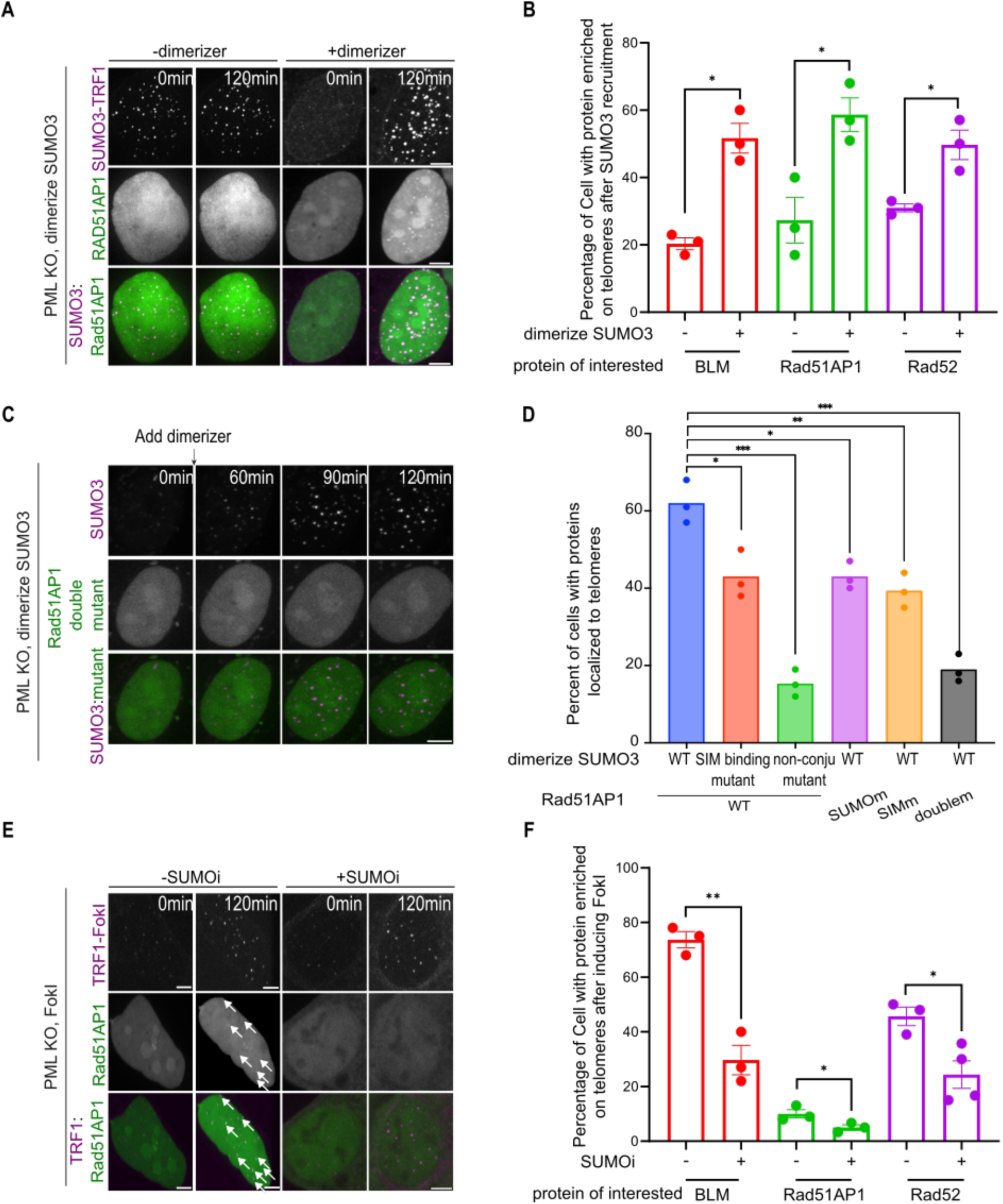
SUMO promotes DNA repair factor enrichment at telomeres independent of APBs. **(A)** Representative images of PML KO cells expressing mCherry-eDHFR-SUMO3, 3xHalo-TRF1, and GFP-Rad51AP1 with/without adding dimerizer to dimerize SUMO3 to TRF1 at indicated time points. **(B)** Quantification of the percentage of cells that have indicated proteins enriched on telomeres after dimerizing SUMO3 to PML KO telomeres for 2 hours (each dot represents one independent experiment, more than 15 cells per group, three independent experiments). **(C)** Representative images of PML KO cells expressing mCherry-eDHFR-SUMO3, 3xHalo-TRF1, and GFP-Rad51AP1 SUMO/SIM double mutant after adding dimerizer at the indicated time point. **(D)** Quantification of cells that have indicated proteins enriched to PML KO telomeres for 2 hours of adding dimerizes. **(E)** Representative images of PML KO cells expressing mCh-TRF1-FokI and GFP-Rad51AP1 with the treatment of 4-Hydroxyestradiol (4-OHT) to induce DNA damage at an indicated time point, treated with/without 1 μM SUMOi for 2 days. White arrows indicate Rad51AP1 co-localizations with telomeres. **(F)** Quantification of PML KO cells that have indicated proteins enriched on telomeres after inducing FokI-TRF1 and with/without 1 μM SUMOi treatment for 2 days. Scale bars, 5 μm.

Significantly, sumoylation per se is required for the enrichment of BLM, Rad51AP1, and Rad52 at telomeres, since much less enrichment was seen when non-conjugatable SUMO1 or SUMO3 was used or when SUMOi was applied (Fig. 3D, Fig. S5C-F). The effect also required the SUMO-SIM interaction, since reduced levels of the three repair proteins were found at telomeres when a SUMO3 mutant defective in SIM binding was used (Fig. 3D, Fig. S5C-H).

Consistently, the enrichment of Rad51AP1 was diminished when its sumoylation and SIM sites were mutated separately or together (Fig. 3C, D). These data support the conclusion that sumoylation and SUMO-SIM interaction contribute to telomere enrichment of BLM, Rad51AP1, and Rad52 in the absence of PML and APBs.

Finally, we used the FokI-induced telomere break system described above in PML null U2OS cells to assess the contribution of sumoylation to telomeric localization of BLM, Rad51AP1, and Rad52 in ALT cells. After induction with FokI, but not a FokI enzymatically dead mutant, BLM, Rad52, and Rad51AP1 were found to be localized at telomeres (Fig. 3F, Fig. S6C), suggesting that their recruitment to ALT telomeres is DNA break-dependent and PML-independent. In addition, telomere recruitment of BLM, Rad51AP1, and Rad52 was reduced upon SUMOi treatment (Fig. 3E, F, Fig. S6A, B). This result is consistent with the notion that SUMO contributes to the recruitment of the three repair factors to ALT telomeres depending on DNA damage response but independent of PML and APBs.

### Rad52 recruitment induces SUMO enrichment and signatures of phase separation

Given that SUMO enriches BLM, Rad52, and Rad51AP1 at telomeres to promote ALT, we asked whether directly targeting BLM, Rad52, and Rad51AP1 at telomeres can promote ALT features. To this end, we directed Rad52, Rad51AP1, and BLM individually to telomeres in PML null U2OS cells using the dimerizer system described above (Fig. 4A). Live cell imaging of the GFP-TRF1 foci, which represent telomeres, showed increase in intensity and decrease in numbers over time, suggesting the formation of phase-separated condensates containing telomeres (Fig. 4B, Fig. S7A-E, Movie 7-9). Importantly, all three tested proteins formed bright and round foci that showed fusion behaviors characteristic of phase-separated condensate (Fig. 4B, Fig. S7A-E, Movie 7-9). As a result of condensate fusion in all three cases, telomeres became clustered, as confirmed by telomeric DNA FISH (Fig. S7J). These effects are reminiscent of those seen upon telomere targeting SUMO or SIM described above (Fig. 2E). Thus, telomere-targetting BLM, Rad52, and Rad51AP1 in PML null cells can induce nuclear structures with features of phase-separated condensates that contain both telomeres and DNA repair proteins, reminiscent of APBs.

**Fig. 4.**
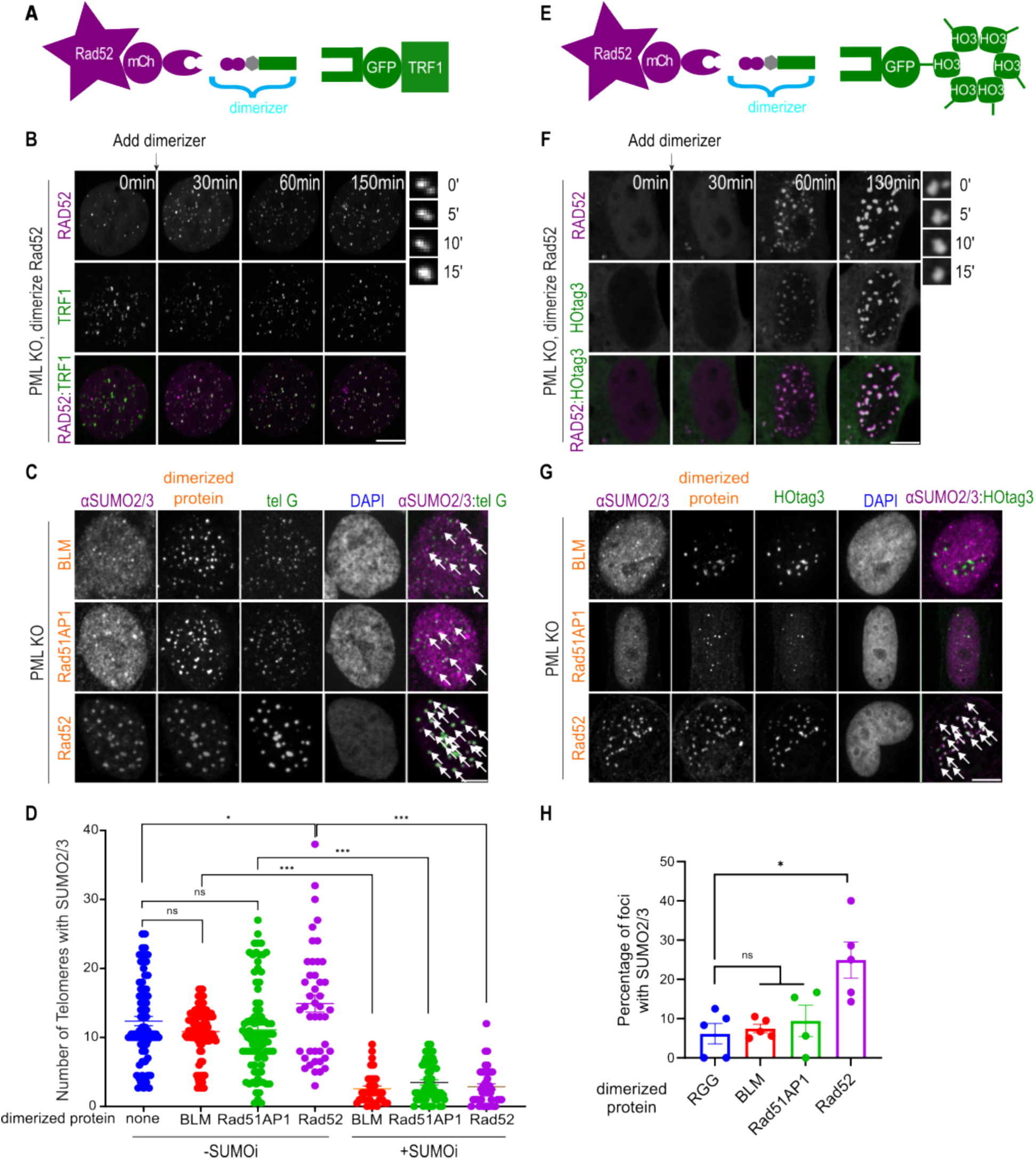
Rad52 recruitmentinduces phase separation and enriches SUMO. **(A)** Dimerization schematic: Rad52/BLM/Rad51AP1 is fused to mCherry and eDHFR, and TRF1 is fused to GFP and 3xHalo enzyme. **(B)** Representative images of PML KO cells after dimerizing mCh-eDHFR-Rad52 to 3xHalo-GFP-TRF1 at indicated time points. Zoomed-in images on the right show a fusion event of TRF1 foci. **(C)** Representative images and **(D)** quantification of SUMO2/3 localization on telomeres in PML KO cells expressing mCh-eDHFR-BLM/Rad51AP1/Rad52 and 3xHalo-TRF1 after adding the dimerizer for 6 hours, with or without 1 μM SUMO inhibitor for 2 days. White arrows indicate SUMO2/3 co-localization on telomeres. **(E)** Dimerization schematic: Rad52/BLM/Rad51AP1 is fused to mCherry and eDHFR, and HOtag3 is fused to GFP and 3xHalo enzyme. **(F)** Representative images of PML KO cells expressing mCh-eDHFR-Rad52 and 3xHalo-GFP-HOtag3 after adding the dimerizer to induce dimerization at indicated time points. Zoomed-in images on the right show a fusion event over time. **(G)** Representative images and **(H)** quantification of SUMO2/3 localization in foci in PML KO cells after dimerizing mCh-eDHFR-BLM, Rad51AP1, Rad52 or RGG to 3xHalo-GFP-HOtag3 for 3 hours (each dot represents one independent experiment, more than 18 cells per group, three independent experiments). Scale bars: 1 μm for the zoomed images and 5 μm for other images.

Among the three tested proteins described above, only telomere targeting of Rad52 led to increased levels of SUMO at telomeres (Fig. 4C, D, Fig. S7F, I). Further, sumoylation per se is required for Rad52-induced SUMO enrichment since SUMOi abolished SUMO enrichment after Rad52 recruitment (Fig. 4D, Fig. S7G-I). Concomitantly, SUMOi also reduced telomere clustering upon telomere-targeting Rad52 (Fig. S7J). This effect was not seen for BLM or Rad51AP1. These results suggest that Rad52’s effects on telomeres are uniquely connected to sumoylation.

Next, we tested whether Rad52 had the intrinsic ability to phase separate and enrich SUMO or whether the observed phase separation and SUMO enrichment were due to Rad52’s function at the telomere. To this end, we used an established method to test Rad52’s ability to phase separate off telomeres in PML null cells (Lackner et al., 2022; Q. Zhang et al., 2018) (Fig. 4E). In the method, the potential phase separation protein is dimerized to an oligomer, the synthetic hexamer HOtag3, which would increase interaction valence between the phase separation proteins to induce condensation formation. Live imaging showed that before dimerization, both Rad52 and HOtag3 had diffusively localized signals in the nucleoplasm (Fig. 4F, Movie 10). After adding the dimerizer, condensates containing both Rad52 and HOtag3 were formed. In addition, the condensates coarsened over time through coalescence, suggesting liquid properties of the condensates. Since Rad52 in budding yeast has been reported to undergo phase separation to facilitate DNA repair (Oshidari et al., 2020), our observations suggest that the ability of Rad52 to phase separate is conserved.

Similar to Rad52, we found dimerizing Rad51AP1 and BLM to HOtag3 also led to condensate formation (Fig. S8A, B, Movie 11-12). Phase diagram mapping indicated that BLM had a higher, while Rad51AP1 had a lower, propensity to phase separate than Rad52 (Fig. S8C). However, SUMO2/3 enrichment, as shown by immunofluorescence, only occurred in the synthetic Rad52 condensates, but not in BLM, Rad51AP1, or another synthetic condensate formed by dimerizing the phase separating arginine/glycine-rich (RGG) domain from the P granule component LAF-1 protein (Elbaum-Garfinkle et al., 2015; Schuster et al., 2018) to HOtag3 (Fig. 4G, H, Fig. S8D). Interestingly, we found SUMO1 was not very significantly enriched in Rad52 condensates (Fig. S8E, F). Together these results suggest that though BLM, Rad52, and Rad51AP1 all have the ability to phase separate, only Rad52 has the intrinsic property to enrich SUMO.

### SUMO promotes Rad52 collaboration with BLM for telomere DNA synthesis

To explore the relationship between Rad52 and SUMO further, we investigated if Rad52 recruitment to telomeres could enrich other DNA repair factors in a SUMO-dependent manner. Live imaging showed that recruiting Rad52 to PML null telomeres enriched BLM and Rad51AP1 (Fig. 5A, B). The enrichment of both proteins, along with SUMO1 and SUMO2/3, was reduced after adding the SUMO inhibitor (Fig. 5A, B). Conversely, recruiting BLM and Rad51AP1 also enriched Rad52 and the enrichment can be reduced by SUMOi treatment (Fig. S9A-C), but the degree of enrichment was much less than BLM and Rad51AP1 enrichment after recruiting Rad52. Consistent with this difference, in PML-containing control cells, recruiting Rad52 led to APB formation, while recruiting Rad51AP1 and BLM did not (Fig. S9D, E). These results suggest that Rad52 has a unique ability to enrich SUMO and DNA repair proteins at telomeres and to promote APB formation in ALT cells.

**Fig. 5.**
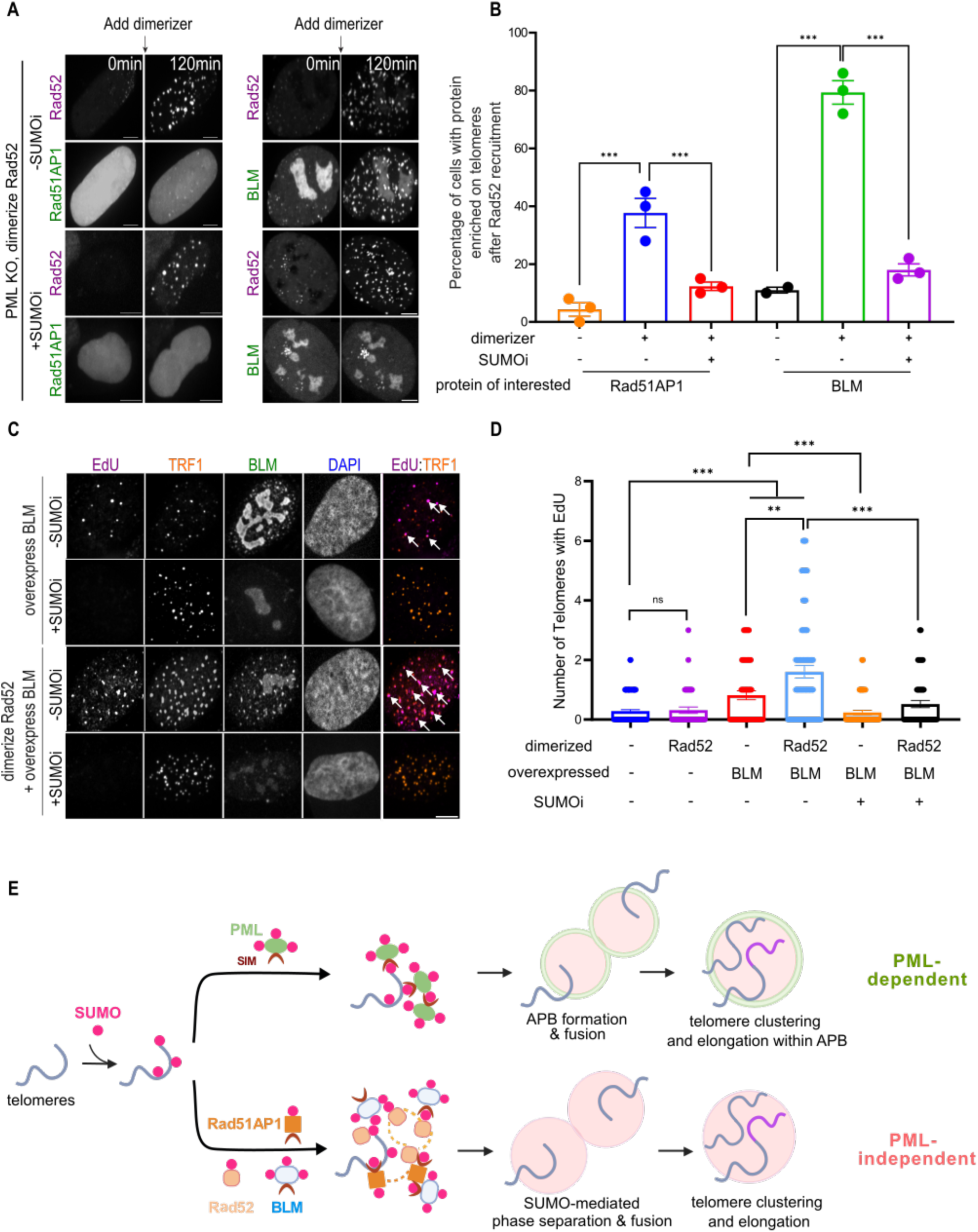
SUMO promotes Rad52 collaboration with BLM for telomere DNA synthesis. **(A)** Representative images and **(B)** quantification of Rad51AP1 and BLM localization on telomeres after Rad52 dimerization in PML KO cells expressing GFP-Rad51AP1 or GFP-BLM after dimerizing mCh-eDHFR-Rad52 to 3xHalo-TRF1, with or without 1 μM SUMOi for 2 days. **(C)** Representative images and **(D)** quantification of EdU assay showing newly synthesizedtelomeric DNA after dimerizing Rad52 to telomeres in PML KO and WT, with or without overexpressing BLM and treating with 1 μM SUMO inhibitor for 2 days. Scale bars, 5 μm. **(E)** Model: in the absence of PML, SUMOylation after DNA damage at ALT telomeres recruits DNA repair factors, including Rad52, Rad51AP1, and BLM, and promotes their co-phase separation for telomere clustering and elongation without APBs.

Next, we examined the effects of telomere targeting Rad52, BLM, and Rad51AP1 on telomere DNA synthesis. We found that Rad52 and BLM, but not Rad51AP1, recruitment to telomeres was sufficient to induce telomere DNA synthesis in PML-containing cells (Fig. S9F, G). However, in PML null cells, recruiting BLM, but not Rad52 and Rad51AP1, to telomeres induced telomere DNA synthesis and SUMOi abolished BLM-induced telomere DNA synthesis (Fig. S9F, G). This finding confirmed the critical role of BLM in ALT telomere DNA synthesis(Loe et al., 2020; Min et al., 2019; J.-M. Zhang et al., 2021), and further suggests that BLM function in ALT can be APB-independent but still requires SUMO. Significantly, Rad52 telomere targeting enhanced telomere synthesis induced by BLM overexpressing in PML null cells, which is abolished by SUMOi treatment, indicating the dependence on sumoylation (Fig. 5C, D). Together, these data suggest that SUMO promotes Rad52 collaboration with BLM to support ALT telomere DNA synthesis beyond APB formation in ALT cells.

## Discussion

Previous studies conducted in PML-containing ALT cells have established an important role of SUMO in ALT via promoting APB formation. Building on the recent establishment of PML null ALT cell lines, we addressed whether SUMO also contributes to ALT in a PML- and APB-independent manner. We provide multiple lines of evidence to support an intrinsic ability of sumoylation in promoting multiple ALT features in PML null cells. We show that this ability is mediated by SUMO-SIM interaction, and we have identified DNA repair proteins involved in ALT that can mediate SUMO-based contribution to ALT features independent of PML. We further unveiled a unique ability of Rad52 to mediate the enrichment of SUMO at telomeres and SUMO-based enabling of telomere clustering, as well as its positive effect on telomere DNA synthesis. Collectively, our work defines the important roles of sumoylation and SUMO-SIM interaction in promoting both branches of the ALT processes independent of PML.

Utilization of a SUMO inhibitor and a chemically induced protein-protein dimerization method, we examined the roles of sumoylation and telomere targeting of SUMO isoforms, SIM, and DNA repair proteins. We queried endogenously ALT features and those induced by telomere breaks. Using both live cell imaging and immunofluorescent imaging, we systematically examined the telomere localization of these proteins, telomere clustering, and telomere DNA synthesis. Our data suggest that sumoylation at ALT telomeres can directly recruit DNA repair proteins via SUMO-SIM interaction and drive telomere clustering and telomere DNA synthesis in the absence of PML and APBs (Fig. 5E). The proteins enriched via SUMO include helicase BLM known to be important for ALT, as well as Rad52 and Rad51AP1, two proteins that control each of the ALT pathways, suggesting SUMO can mediate both branches of the ALT processes. In addition, we discover that, though not required, APBs still play important roles in ALT, and these roles are intricately linked to sumoylation. We thus conclude that sumoylation and not APB formation per se is a fundamental requirement for ALT. These data let us propose that targeting the SUMO pathway using SUMOi can efficiently abolish ALT, thus providing a potentially effective approach for ALT cancer therapy.

Between the two proteins promoting separate ALT pathways, Rad52 has a unique ability to enrich SUMO and enhance telomere clustering and telomere DNA synthesis. These effects are reminiscent of PML, though Rad52 does not have a wide range of functions as PML. What is the unique biochemical nature of Rad52 that renders its unique connection with SUMO in the ALT pathway remains to be determined. We speculate that these features may be related to its ability to interact with the only SUMO E2 enzyme UBC9(Ouyang et al., 2009), an interaction that has not been reported for other proteins involved in ALT. Future work to test this hypothesis and other possible mechanisms to further clarify the roles of Rad52 in ALT would be interesting.

In PML-containing ALT cells, it is known that SUMO mediates PML protein phase separation to form APBs(H. Zhang et al., 2020). We also observed signatures of phase separation after recruiting SUMO and DNA repair factors to telomeres in PML null cells, which might be linked to the intrinsic ability of these DNA repair proteins, including Rad52, BLM, and Rad51AP1, to undergo phase separation. Since SUMO has been established to act as a glue to promote the co-enrichment of repair factors at the DNA damage site(Psakhye & Jentsch, 2012), we suggest a model where SUMO mediates co-phase separation of DNA repair factors at ALT telomeres in the absence of PML (Fig. 5E). It is worth noting that these repair proteins have lower propensity to phase separate than PML because unlike PML, expressing these protein alone in cells does not form condensates. Therefore, even though not required, gaining access to PML protein can enhance SUMO-mediated phase separation of repair factors at ALT telomere for more efficient telomere clustering and telomere DNA synthesis. We suspect that SUMO-mediated co-phase separation of repair factors may be used at non-telomere damage sites to enrich multiple DNA repair proteins and DNA substrates. Future experiments to test these ideas will further clarify how SUMO collaborates with a wide range of DNA repair factors in many DNA repair processes for genome protection.

## Materials and Methods

### Cell culture

All WT experiments were performed with U2OS cells. PML KO cells were gifts from Dr. Eros Lazzerini Denchi. U2OS cells with endogenous PML tagged with Clover (U2OS Clover-PML) are previously described(Pinder et al., 2015). Cells were cultured in growth medium (Dulbecco’s Modified Eagle’s medium with 10% FBS and 1% penicillin–streptomycin) at 37 °C in a humidified atmosphere with 5% CO2.

### Plasmids

The plasmid for inducing DNA damage at telomeres (mCherry-TRF1-FokI) was previously published(Cho et al., 2014). 3xHalo-GFP-TRF1 was previously published(H. Zhang et al., 2020). SIM (or SIM mutant) for mCherry-eDHFR-SIM, PML WT, PML SUMOylation sites, and non-conjugatable SUMOs are from plasmids gifted by Michael Rosen(Banani et al., 2016). SUMO1/2/3 (or SUMO mutant) for mCherry-eDHFR-SUMO is from plasmids gifted by Karsten Rippe(Chung et al., 2011). SUMO SIM interacting mutant was generated by mutating the FKIK (SUMO3) or FKVK (SUMO1) sequence in each SUMO module to FAAA(Banani et al., 2016). Non-conjugatable SUMO modules were generated by mutating the C-terminal di-Gly motif to di-Val(Banani et al., 2016). SIM mutant was generated by mutating the VIDL sequence into VADA(Banani et al., 2016). The vector containing mCherry-eDHFR is from our published plasmid Mad1-mCherry-eDHFR(H. Zhang et al., 2017). NLS was cloned in 3xHalo-GFP-HOTag3(Lackner et al., 2022). RGG-mCherry-RGG-eDHFR was previously published(Lackner et al., 2022). GFP-BLM is from Addgene plasmid #80070. GFP-Rad51AP1 and mutants are gifted by Dr. Roderick J. O’Sullivan and introduced into target plasmids through in-fusion cloning (#638948, Takara Bio). All the target plasmids in this study are derived from a plasmid that contains a CAG promoter for constitutive expression, obtained from E. V. Makeyev(Khandelia et al., 2011).

### Cell treatment with chemicals

SUMO inhibitor ML-792 (HY-108702, Selleck Chem) used to inhibit SUMOylation was added to cells at 1 μM for 2 days. For triggering telomere DNA damage in FokI cells, 4-hydroxytamoxifen (4-OHT) (H7904, Sigma-Aldrich) was added to cells at 1 μM working concentration for the indicated time.

### Synchronize cells to G2

Cells were first treated with 2 mM thymidine (T1895, Sigma-Aldrich) for 21 h, released into fresh medium for 4 h, and then treated with 15 μM CDK1i (RO-3306, cat# SML0569, Sigma-Aldrich) for 12 h.

### Protein dimerization with chemical dimerizers

Design, synthesis, and storage of the dimerizer TFH (**T**MP-**F**luorobenzamide-**H**alo) were reported previously(Lackner et al., 2022). Dimerization on telomeres was performed as previously described(Xu et al., 2022). Briefly, TFH was added directly to the growth medium to a final working concentration of 100 nM. Cells were incubated with the dimerizer-containing medium for the indicated times, followed by immunofluorescence (IF) or fluorescence in situ hybridization (FISH). For live imaging with protein dimerization, the dimerizers are first diluted to 200 nM in growth medium and then further added to cell chambers to the working concentration after first-round imaging.

### Immunofluorescence (IF) and telomere DNA fluorescence in situ Hybridization (FISH)

IF and FISH were performed following a previously published protocol(Zhao et al., 2021). Cells were fixed in 4% formaldehyde for 10 min at room temperature, followed by permeabilization in 0.5% Triton X-100 for 10 min. Cells were incubated with primary antibody at 4°C in a humidified chamber overnight and then with secondary antibody for 1 h at room temperature before washing and mounting. Primary antibodies were anti-SUMO1 (Ab32058, Abcam,1:200 dilution), anti-SUMO2/3 (Asm23, Cytoskeleton, 1:200 dilution), and anti-PML (sc966, Santa Cruz, 1:50 dilution). For IF-FISH, coverslips were first stained with primary and secondary antibodies, then fixed again in 4% formaldehyde for 10 min at room temperature. Coverslips were then dehydrated in an ethanol series (70%, 80%, 90%, 2 min each) and incubated with a 488-tel G or Cy3-tel C PNA probe (F1008, F1002, Panagene) at 75°C for 5 min and then overnight in a humidified chamber at room temperature. Coverslips were then washed and mounted for imaging.

### Telomere DNA synthesis detection by EdU

Following transfection, cells were pulsed with EdU (10 μM) along with protein dimerization or DNA damage induction for 6 hrs before harvest. Cells on glass coverslips were washed twice in PBS and fixed with 4% paraformaldehyde (PFA) for 10 min. Cells were permeabilized with 0.3% (v/v) Triton X-100 for 5 min. The Click-IT Plus EdU Cell Proliferation Kit with Alexa Flour 488 and 647 (C10633, C10635 Invitrogen) was applied to cells for 30 minutes to detect EdU.

### Cell imaging and image processing

Imaging acquisition was performed as previously described(Xu et al., 2022). For live imaging, cells were seeded on 22 x 22 mm glass coverslips coated with poly-D-lysine (P1024, Sigma-Aldrich). When ready for imaging, coverslips were mounted in magnetic chambers (Chamlide CM-S22-1, LCI) with cells maintained in a normal medium supplemented with 10% FBS and 1% penicillin/streptomycin at 37 °C on a heated stage in an environmental chamber (TOKAI HIT Co., Ltd.). Images were acquired with a microscope (ECLIPSE Ti2) with a 100 × 1.4 NA objective, a 16 XY Piezo-Z stage (Nikon Instruments Inc.), a spinning disk (Yokogawa), an electron multiplier charge-coupled device camera (IXON-L-897), and a laser merge module that was equipped with 488 nm, 561 nm, 594 nm, and 630 nm lasers controlled by NIS-Elements Advanced Research. For both fixed cells and live imaging, images were taken with 0.5 μm spacing between Z slices, for a total of 8 μm. For movies, images were taken at 5 min intervals for up to 3 hr.

Images were processed and analyzed using NIS-Elements AR (Nikon). Maximum projections were created from z stacks, and thresholds were applied to the resulting 2D images to segment and identify telomere/ SUMO foci as binaries. For colocalization quantification of two fluorescent labels, fixed images were analyzed by the binary operation in NIS-Elements AR. Colocalized foci were counted if the objects from different layers contained overlapping pixels.

### Statistical methods

All error bars represent means ± SEM. Statistical analyses were performed using Prism 10.0 (GraphPad software). Two-tailed nonpaired t-tests have been used for all tests. Statistical significance: N.S., not significant, P > 0.05; *P < 0.05; **P < 0.01; ***P < 0.001.

## Acknowledgments

We thank Dr. Roger A. Greenberg (University of Pennsylvanian), Dr. Eros Lazzerini Denchi (National Cancer Institute), Dr. Michael Rosen (UT Southwestern), Dr. Karsten Rippe (German Cancer Research Center) for kindly gifting plasmids and cell lines. This work was supported by the United States National Institutes of Health Grant U01CA260851 to HZ, GM118510 to DC, R35 GM145260 to XZ., a Project Grant from the Canadian Institutes of Health Research (CIHR) PJT-156017 to GD. A.R.W. receives support from NIH training awardsT32 GM133332 (Dept. P&CB, University of Pittsburgh) and Ruth L. Kirschstein National Research Scientist Training Award CA278287.

## Author contributions

H.Z., R.Z., M.X., and X.Z. conceptualized this study. R.Z. designed and conducted the experiments. R. L. and D.C. designed and synthesized the dimerizers. J. S. and G. D. made the U2OS PML-Clover cell line. R.Z., A.R.W., H.Z., X.Z. and R. O. S. analyzed the results. R.Z. made the figures. R.Z., H.Z., and X.Z. wrote the manuscript with comments from other authors.

## Data availability

All data needed to evaluate the conclusions in the paper are present in the paper and the Supplementary Materials.

---

Supplementary Materials for

**This file includes:**

Supplementary Figures 1 to 9; Supplementary Movies 1 to 12

**Movie 1** Inducing DNA damage at PML KO telomeres. Movie for Fig. 1D. 4-OHT was added to cells after the first time point to induce damage. The box shows a telomere fusion event. Scale bars, 5 μm.

**Movie 2** Inducing DNA damage at PML-Clover telomeres. Movie for Fig. 1E. Left: Composite of FokI-TRF1 (magenta) and PML (green), middle: mCh-FokI-TRF1, right: PML. 4-OHT was added to cells after the first time point to induce damage. The boxes show telomere fusion without forming APBs. Scale bars, 5 μm.

**Movie 3** Dimerizing SUMO3 to PML KO telomeres. Movie for Fig. 2B. Left: Composite of SUMO3 (magenta) and TRF1 (green), middle: mCh-eDHFR-SUMO3, right: 3xHalo-GFP-TRF1. The dimerizer was added to cells after the first time point to induce dimerization. The box shows a telomere fusion event. Scale bars, 5 μm.

**Movie 4** Dimerizing SUMO1 to PML KO telomeres. Movie for Fig. S3A. Left: Composite of SUMO1 (magenta) and TRF1 (green), middle: mCh-eDHFR-SUMO1, right: 3xHalo-GFP-TRF1. The dimerizer was added to cells after the first time point to induce dimerization. The box shows a telomere fusion event. Scale bars, 5 μm.

**Movie 5** Dimerizing SUMO2 to PML KO telomeres. Movie for Fig. S3A. Left: Composite of SUMO2 (magenta) and TRF1 (green), middle: mCh-eDHFR-SUMO2, right: 3xHalo-GFP-TRF1. The dimerizer was added to cells after the first time point to induce dimerization. The box shows a telomere fusion event. Scale bars, 5 μm.

**Movie 6** Dimerizing SUMO3 SIM interacting mutant to PML KO telomeres. Movie for Fig. 2B. Left: Composite of SUMO3 mutant (magenta) and TRF1 (green), middle: mCh-eDHFR-SUMO3m, right: 3xHalo-GFP-TRF1. The dimerizer was added to cells after the first time point to induce dimerization. Scale bars, 5 μm.

**Movie 7** Dimerizing Rad52 to PML KO telomeres. Movie for Fig. 4B. Left: Composite of Rad52 (magenta) and TRF1 (green), middle: mCh-eDHFR-Rad52, right: 3xHalo-GFP-TRF1. The dimerizer was added to cells after the first time point to induce dimerization. The box shows a telomere fusion event. Scale bars, 5 μm.

**Movie 8** Dimerizing BLM to PML KO telomeres. Movie for Fig. S7A. Left: Composite of BLM (magenta) and TRF1 (green), middle: mCh-eDHFR-BLM, right: 3xHalo-GFP-TRF1. The dimerizer was added to cells after the first time point to induce dimerization. The box shows a telomere fusion event. Scale bars, 5 μm.

**Movie 9** Dimerizing Rad51AP1 to PML KO telomeres. Movie for Fig. S7C. Left: Composite of Rad51AP1 (magenta) and TRF1 (green), middle: mCh-eDHFR-Rad51AP1, right: 3xHalo-GFP-TRF1. The dimerizer was added to cells after the first time point to induce dimerization. The box shows a telomere fusion event. Scale bars, 5 μm.

**Movie 10** Dimerizing Rad52 to HOtag3 in PML KO cells. Movie for Fig. 4F. Left: Composite of Rad52 (magenta) and HOtag3 (green), middle: mCh-eDHFR-Rad52, right: 3xHalo-GFP-HOtag3. The dimerizer was added to cells after the first time point to induce dimerization. The box shows a droplet fusion event. Scale bars, 5 μm.

**Movie 11** Dimerizing BLM to HOtag3 in PML KO cells. Movie for Fig. S8A. Left: Composite of BLM (magenta) and HOtag3 (green), middle: mCh-eDHFR-BLM, right: 3xHalo-GFP-HOtag3. The dimerizer was added to cells after the first time point to induce dimerization. The box shows a droplet fusion event. Scale bars, 5 μm.

**Movie 12** Dimerizing Rad51AP1 to HOtag3 in PML KO cells. Movie for Fig. S8B. Left: Composite of Rad51AP1 (magenta) and HOtag3 (green), middle: mCh-eDHFR-Rad51AP1, right: 3xHalo-GFP-HOtag3. The dimerizer was added to cells after the first time point to induce dimerization. Scale bars, 5 μm.

**Fig. S1.**
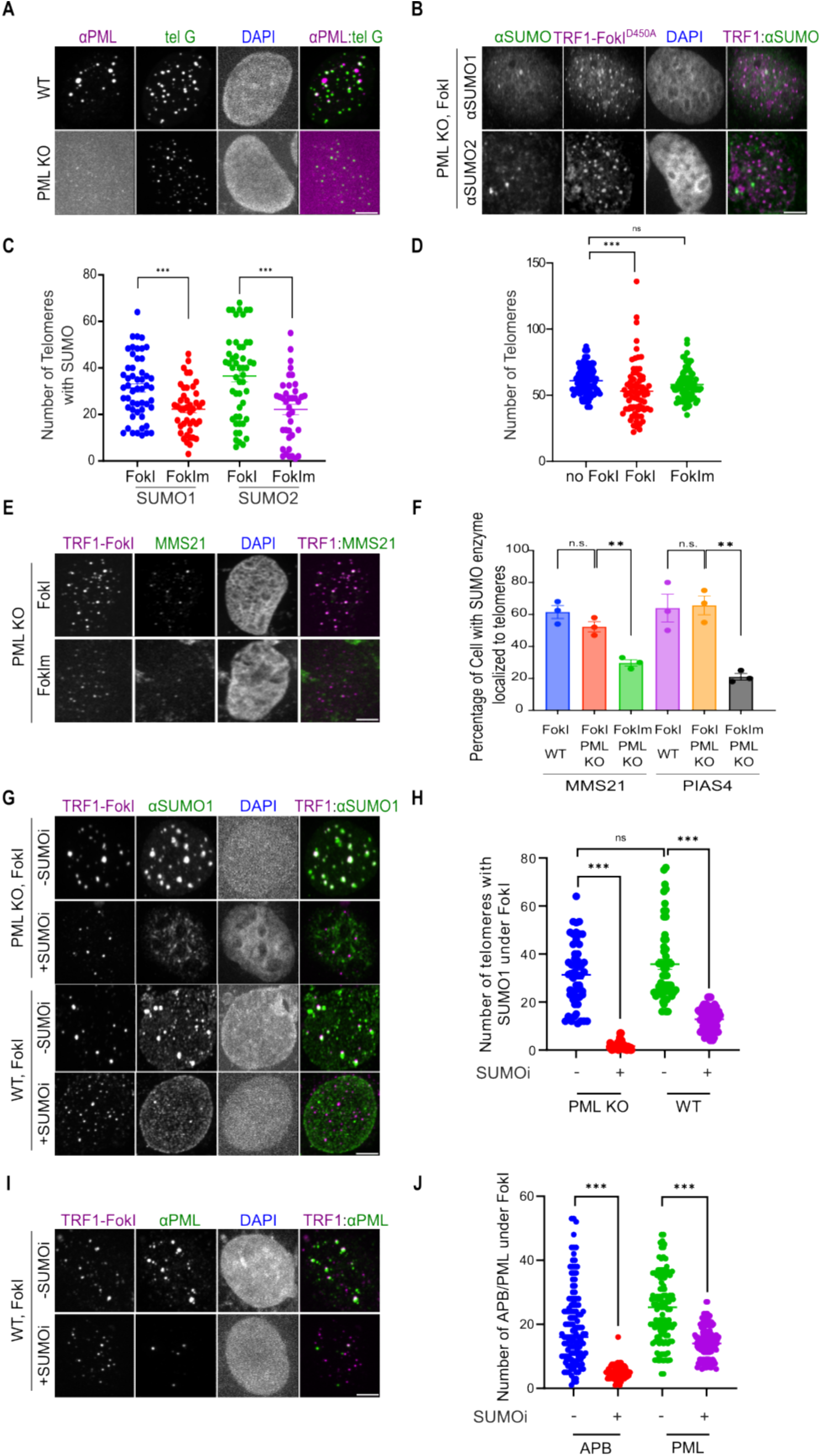
ALT features after inducing telomere-specific double-strand break with/without SUMOi in U2OS WT and PML KO cells. **(A)** Representative images of PML staining in WT and PML KO cells. **(B)** Representative images of SUMO1/2/3 localization on telomeres in PML KO expressing TRF1-FokI-D450A after adding 4-Hydroxyestradiol (4-OHT) for 6 hours. **(C)** Quantification of SUMO1/2/3 localization on telomeres and **(D)** telomere numbers in PML KO with or without expressing TRF1-FokI-WT and TRF1-FokI-D450A to induce DNA damage for 6 hours. **(E)** Representative images and **(F)** quantification of MMS21 and PIAS4 localization on telomeres in U2OS WT and PML KO cells expressing mCh-TRF1-FokI and FokI enzymatic dead mutant TRF1-FokI-D450A with treatment of 4-Hydroxyestradiol (4-OHT) for 6 hours. **(G)** Representative images and **(H)** Quantification of SUMO1 localization on telomeres in PML KO and WT cells with or without 1 μM SUMOi under 6-hour FokI-induced DNA damage. **(I)** Representative images, **(J)** quantification of APB and PML body number in WT treating with or without 1 μM SUMOi under FokI-induced DNA damage for 6 hours. Scale bars, 5 μm.

**Fig. S2.**
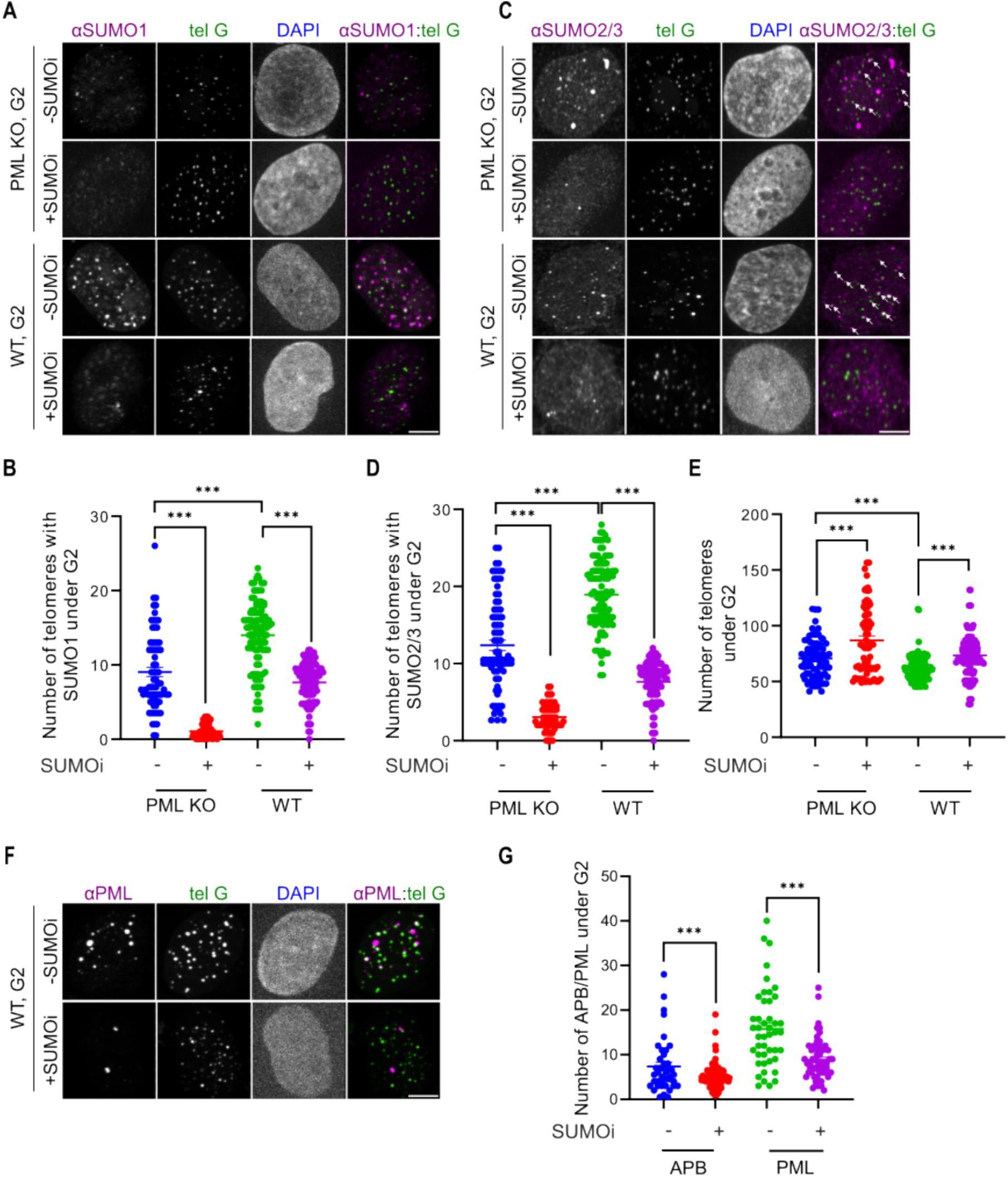
ALT features in G2 arrested-WT and PML KO cells with/without SUMOi treatment. **(A)** Representative images and **(B)** quantification of SUMO1 localization on telomeres in G2 arrested-WT and PML KO cells with or without 1 μM SUMOi after 2 days. **(C)** Representative images and **(D)** quantification of SUMO2/3 localization on telomeres in G2 arrested-WT and PML KO treating with or without 1 μM SUMOi after 2 days. **(E)** Quantification of telomere number in G2 arrested-WT and PML KO treating with or without 1 μM SUMOi after 2 days. **(F)** Representative images, **(G)** quantification of APB and PML body number in G2 arrested-WT treating with or without 1 μM SUMOi after 2 days. Scale bars, 5 μm.

**Fig. S3.**
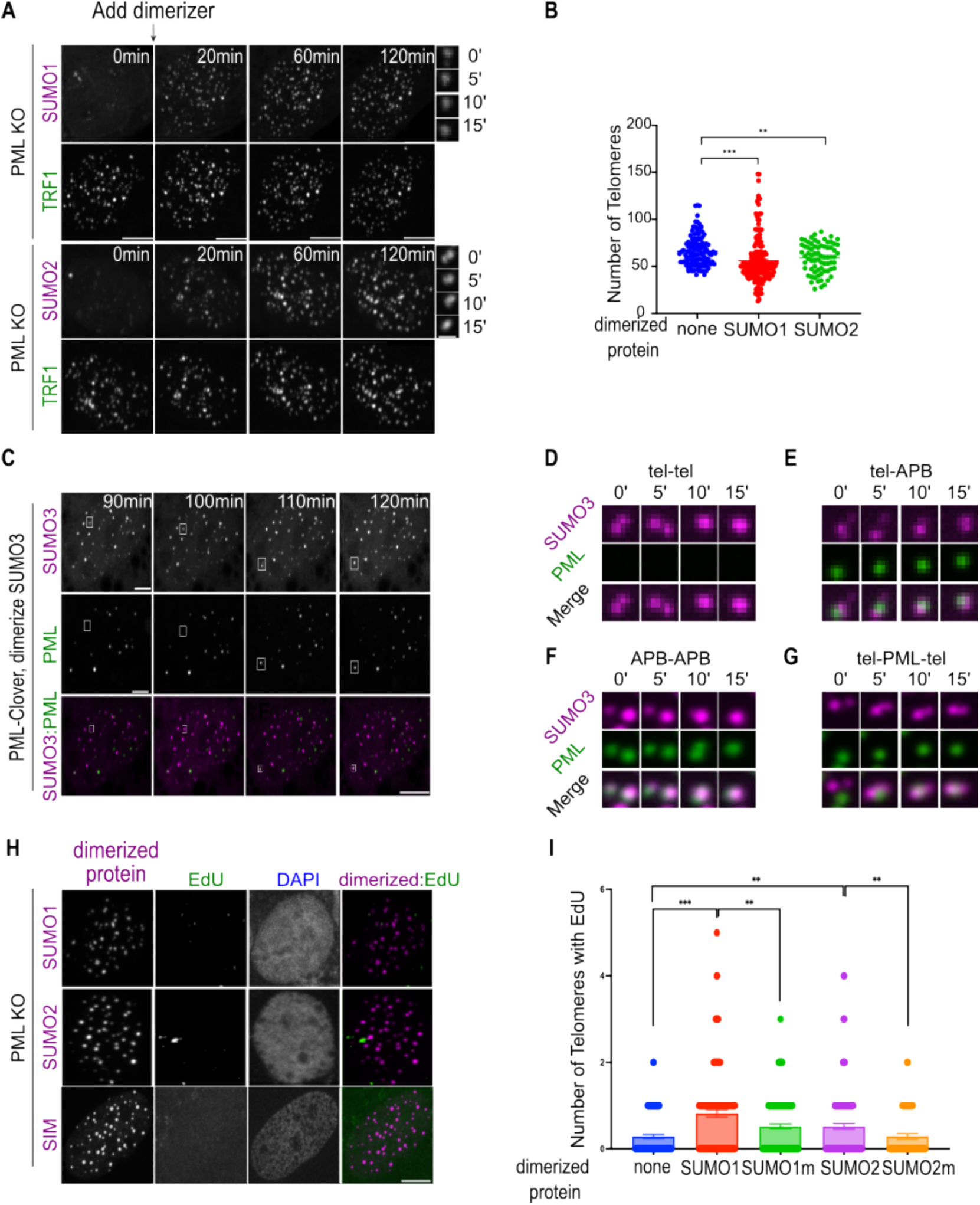
ALT features after dimerizing SUMO to telomeres in PML KO cells. **(A)** Representative images of PML KO cells after dimerizing mCh-eDHFR-SUMO1/SUMO2 to 3xHalo-GFP-TRF1 after the first time point. **(B)** Telomere number after dimerizing mCh-eDHFR-SUMO1/2 to 3xHalo-TRF1 in PML KO cells (more than 50 cells per group, three independent experiments, two-tailed unpaired-test). **(C)** Representative images of PML-Clover cells after dimerizing mCh-eDHFR-SUMO3 to 3xHalo-TRF1 at indicated time points. **(D)-(F)** Zoomed-in images from (C) showing telomere clustering events. **(G)** Zoomed-in images show two telomeres sticking to a PML body. **(H)** Representative images and **(I)** quantification of EdU assay showing newly synthesized telomeric DNA without dimerizing any protein or dimerizing SUMO1/2 and their SIM interaction mutants to telomeres for 6 hours in PML KO cells. Scale bars, 5 μm or 1 μm for the zoomed-in images.

**Fig. S4.**
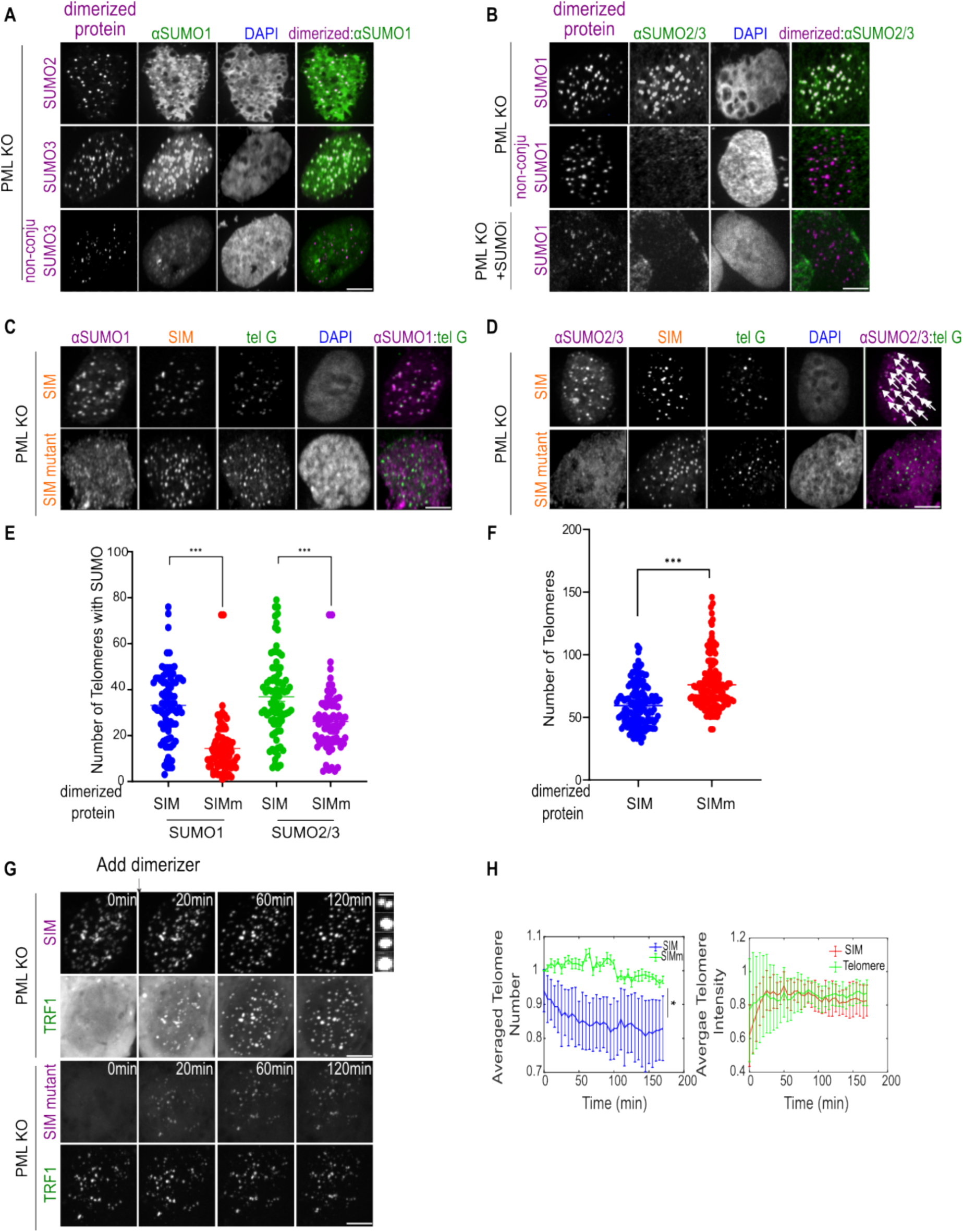
ALT phenotypes after dimerizing SIM and SUMO mutants to telomeres in PML KO cells. **(A)** Immunofluorescent images of SUMO1 in PML KO cells dimerizing mCh-eDHFR-SUMO2/3, non-conjugatable SUMO3 to telomeres for 6 hours. **(B)** Immunofluorescent images of SUMO2/3 in PML KO cells after dimerizing mCh-eDHFR-SUMO1 and non-conjugatable SUMO1 to telomeres for 6 hours, with/without SUMOi. **(C)(D)** Immunofluorescent images and **(E)** quantification of SUMO1/2/3 localization on telomeres and **(F)** telomere numbers in PML KO after dimerizing SIM or SIM mutant to telomeres for 6 hours. **(G)** Representative images of PML KO cells after dimerizing mCh-eDHFR-SIM, or SIM mutant to 3xHalo-GFP-TRF1 at indicated time points. Zoomed-in images show a fusion event of TRF1 foci. **(H)** Telomere number, telomere sum intensity per cell after dimerizing SIM or SIM mutant to PML KO telomeres (more than 21 cells per group, three independent experiments, two-tailed unpaired t-test). Scale bars, 5 μm or 1 μm for zoomed-in images.

**Fig. S5.**
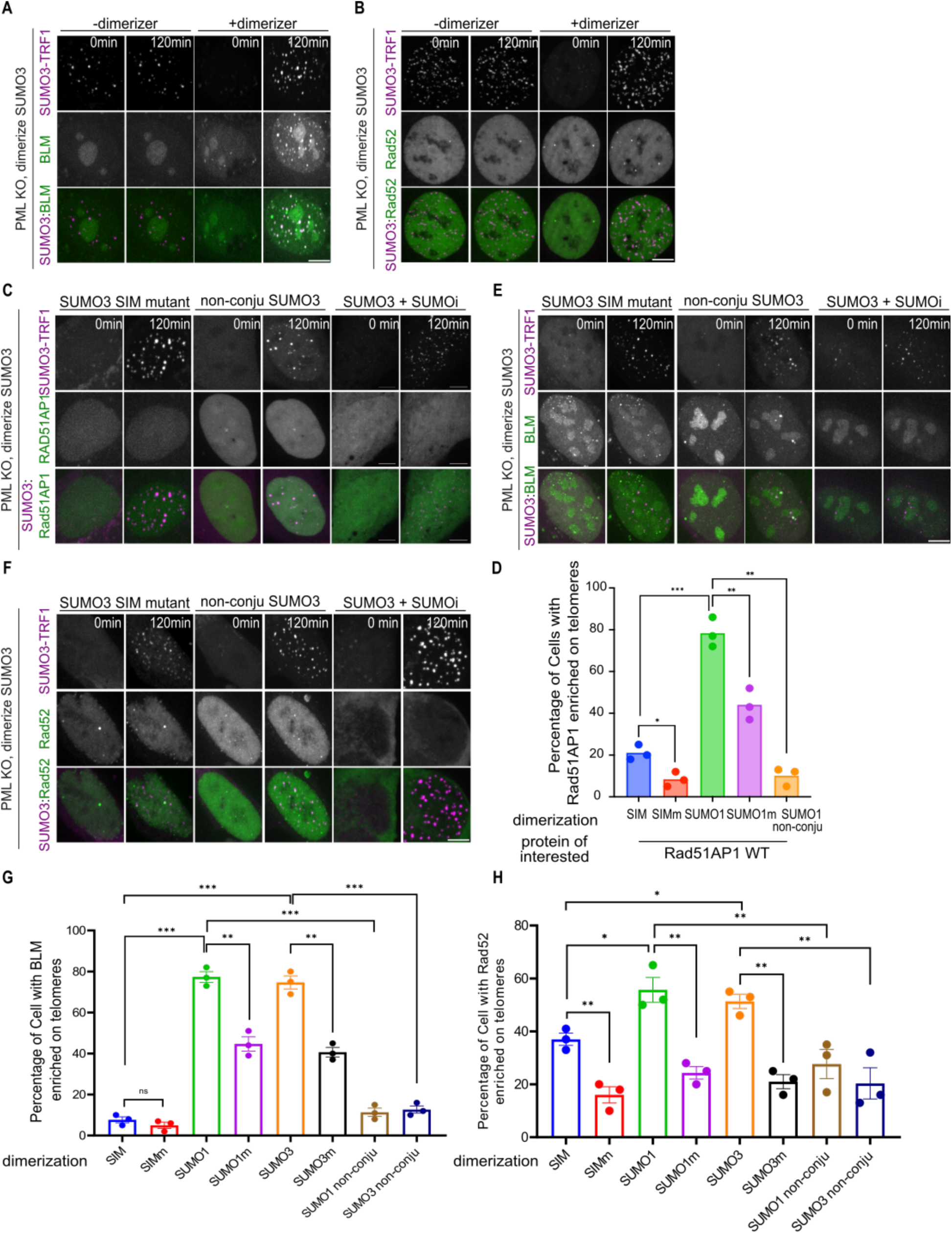
Localization of DNA repair factors to telomeres after dimerizing SUMO to PML KO telomeres. **(A)** Representative images of BLM and **(B)** Rad52 localization on telomeres after dimerizing SUMO3 to telomeres in PML KO cells. **(C)** Representative images of PML KO cells expressing GFP-Rad51AP1, or **(E)** BLM, or **(F)** Rad52 after dimerizing mCh-eDHFR-SUMO3/SIM-binding mutant/non-conjugable mutant to 3xHalo-TRF1, with/without SUMOi at 1 μM for 2 days. **(D)** Quantification of cells with Rad51AP1 or its indicated mutants localized to telomeres after dimerizing SUMO/SIM mutant to PML KO telomeres for 2 hours. **(G)** Quantification of cells that have BLM and **(H)** Rad52 localized to telomeres after dimerizing SUMO/SIM mutant to PML KO telomeres for 2 hours. Scale bars, 5 μm.

**Fig. S6.**
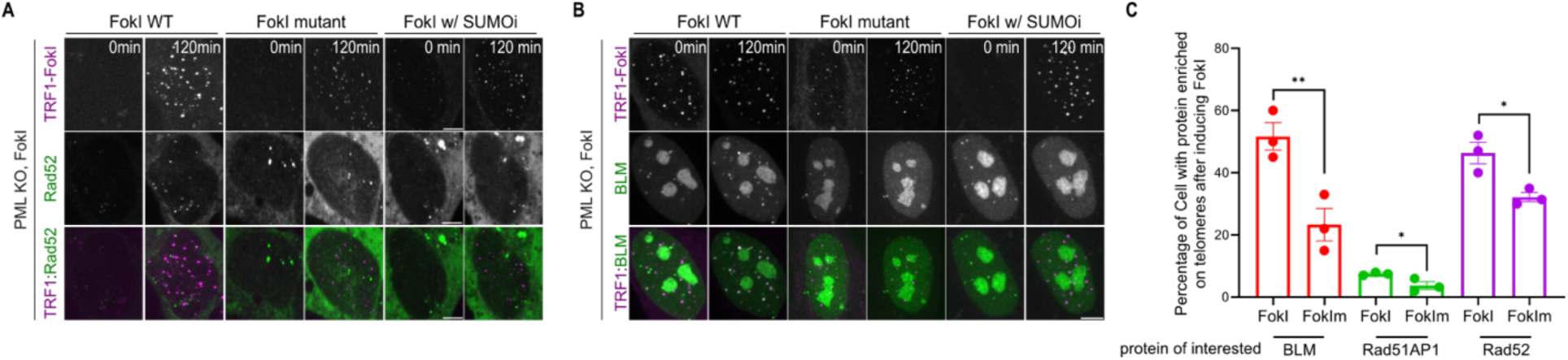
Localization of DNA repair factors to telomeres after inducing FokI in PML KO telomeres. **(A)** Representative images of PML KO cells expressing mCh-TRF1-FokI or enzymatically dead mutant TRF1-FokI-D450A and GFP-Rad52, **(B)** BLM, with the treatment of 4-Hydroxyestradiol (4-OHT) to induce DNA damage at indicated time points, with or without 1 μM SUMOi for 2 days. **(C)** Quantification of cells that have indicated proteins localized to telomeres after inducing DNA damage on PML KO telomeres for 6 hours. Scale bars, 5 μm.

**Fig. S7.**
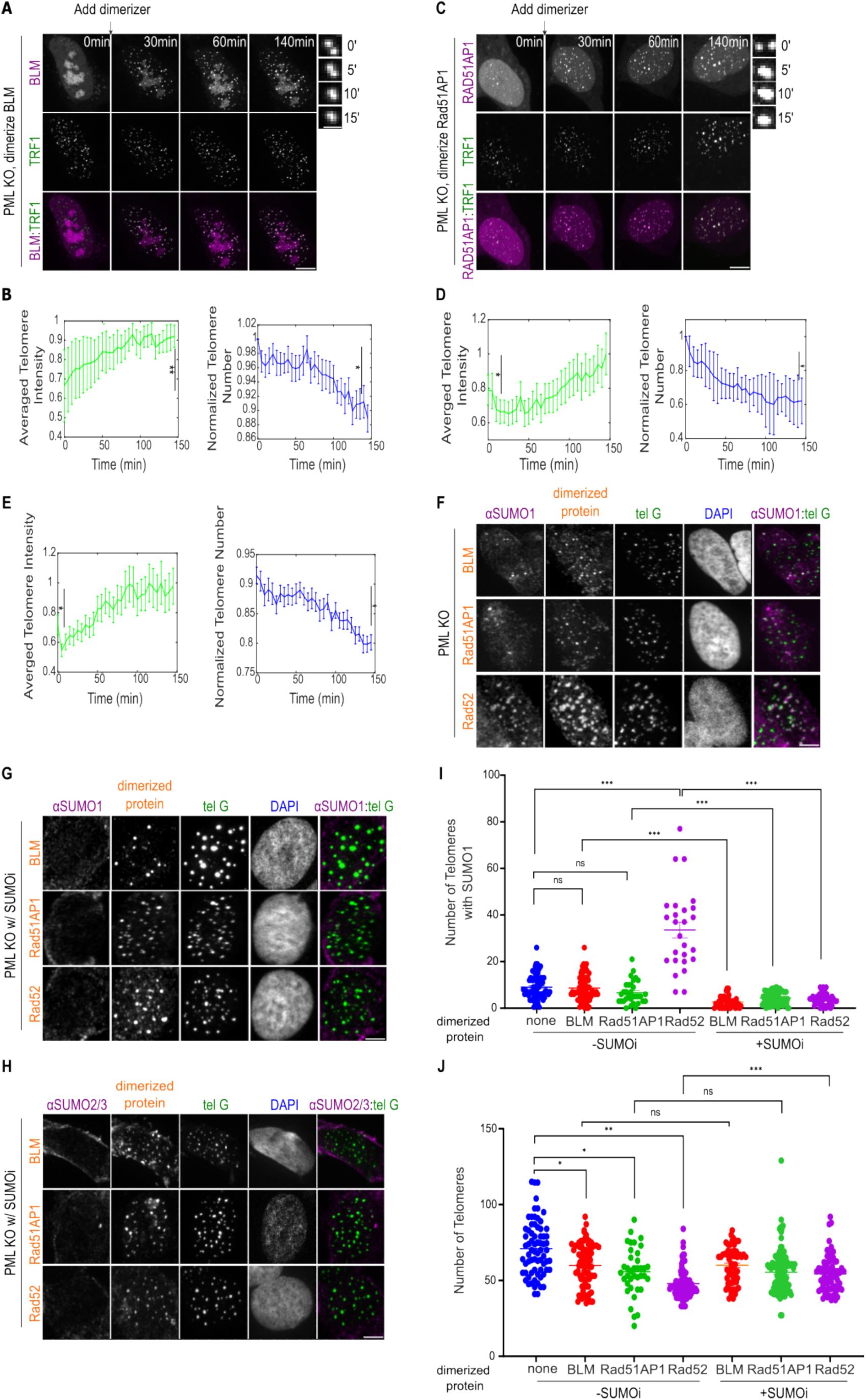
SUMO enrichment after dimerizing DNA repair factors to telomeres in PML KO cells. **(A)** Representative images of PML KO cells after dimerizing mCh-eDHFR-BLM to 3xHalo-GFP-TRF1 at indicated time points. Zoomed-in images show a fusion event of TRF1 foci. **(B)** Telomere sum intensity and telomere number per cell after adding the dimerizer (telomere numbers are normalized by the number at the first time point for each cell, more than 23 cells per group, three independent experiments, two-tailed unpaired *t-test*). **(C)** Representative images of PML KO cells after dimerizing mCh-eDHFR-Rad51AP1 to 3xHalo-GFP-TRF1 at indicated time points. Zoomed-in images show a fusion event of TRF1 foci. **(D)** Telomere sum intensity and telomere number per cell after adding the dimerizer. **(E)** Telomere sum intensity and telomere number per cell after dimerizing mCh-eDHFR-Rad52 to 3xHalo-GFP-TRF1. **(F)** Representative images of SUMO1 localization on telomeres in PML KO cells after dimerizing mCh-eDHFR-BLM/Rad51AP1/Rad52 to 3xHalo-TRF1. **(G)** Representative images of SUMO1 and **(H)** SUMO2/3 localization on telomeres and **(I)** quantification in PML KO cells after dimerizing mCh-eDHFR-BLM/Rad51AP1/Rad52 to 3xHalo-TRF1 with 1 μM SUMOi for 2 days. **(J)** Number of telomeres in PML KO after dimerizing Rad52/BLM/Rad51AP1 to telomeres for 6 hours, with or without 1 μM SUMO inhibitor for 2 days. Scale bars, 5 μm or 1 μm for the zoomed-in images.

**Fig. S8.**
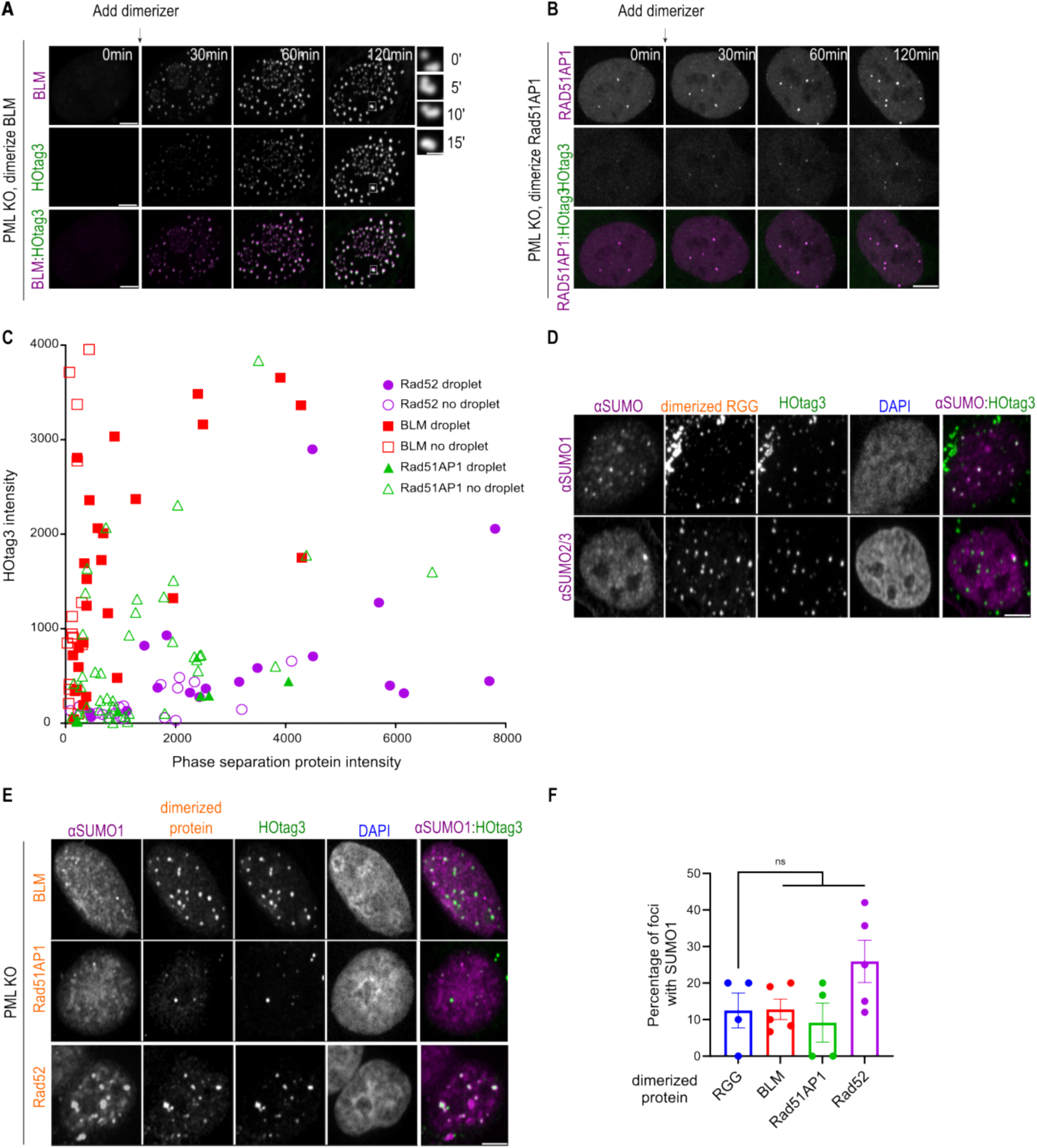
Condensate formation and SUMO enrichment after dimerizing DNA repair factors to HOtag3 in PML KO cells. **(A)** Representative images of PML KO cells after dimerizing mCh-eDHFR-BLM to 3xHalo-GFP-HOtag3 at indicated time points. Inset Zoomed-in images show a fusion event. **(B)** Representative images of PML KO cells after dimerizing mCh-eDHFR-Rad51AP1 to 3xHalo-GFP-HOtag3 at indicated time points. **(C)** Phase diagram of BLM/Rad51AP1/Rad52 droplet formation. Intensities are the mean intensity in cells before dimerization. **(D)** Representative images of SUMO1/2/3 localization in foci in PML KO cells after dimerizing RGG-mCh-eDHFR-RGG to 3xHalo-GFP-HOtag3. **(E)** Representative images and **(F)** quantification of SUMO1 localization in foci in PML KO cells after dimerizing mCh-eDHFR-BLM/Rad51AP1/Rad52 to 3xHalo-GFP-HOtag3. Scale bars, 5 μm or 1 μm for the zoomed-in images.

**Fig. S9.**
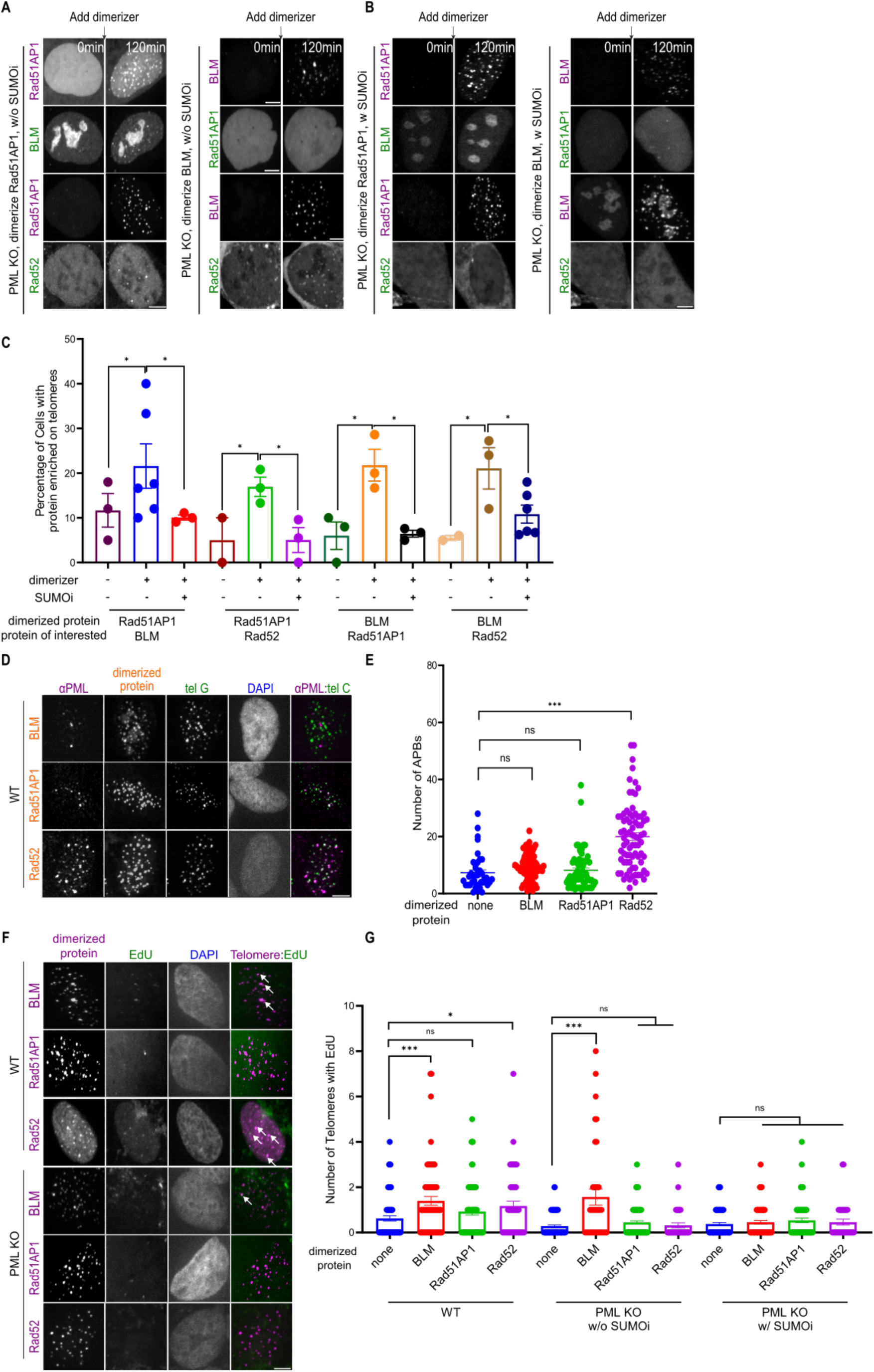
Importance of SUMO for mutual enrichment of repair factors and telomere DNA synthesis. **(A)(B)** Representative images and **(C)** quantification of protein localization to telomeres in PML KO cells expressing GFP-BLM/Rad51AP1/Rad52 after dimerizing mCh-eDHFR-Rad51AP1/BLM to 3xHalo-TRF1, with or without 1 μM SUMOi for 2 days. **(D)** Representative images and **(E)** quantification of APB numbers in WT cells after dimerizing BLM/Rad51AP1/Rad52 to telomeres for 6 hours. **(F)** Representative images of EdU on telomeres after dimerizing BLM, Rad51AP1, or Rad52 to telomeres in PML KO and WT cells. **(G)** Quantification of EdU on telomeres after dimerizing BLM, Rad51AP1, or Rad52 to telomeres in PML KO and WT U2OS cells with or without 1 μM SUMOi for 2 days. Scale bars, 5 μm.

